# Phase dependence of response curves to stimulation and their relationship: from a Wilson-Cowan model to essential tremor patient data

**DOI:** 10.1101/535880

**Authors:** Benoit Duchet, Gihan Weerasinghe, Hayriye Cagnan, Peter Brown, Christian Bick, Rafal Bogacz

## Abstract

Essential tremor manifests predominantly as a tremor of the upper limbs. One therapy option is high-frequency deep brain stimulation, which continuously delivers electrical stimulation to the ventral intermediate nucleus of the thalamus at about 130 Hz. Constant stimulation can lead to side effects, it is therefore desirable to find ways to stimulate less while maintaining clinical efficacy. One strategy, phase-locked deep brain stimulation, consists of stimulating according to the phase of the tremor. In this study, we aim to reproduce the phase dependent effects of stimulation seen in patient data with a biologically inspired Wilson-Cowan model. To this end, we first analyse patient data, and consistently identify in half of the datasets significant dependence of the effects of stimulation on the phase at which stimulation is provided. We approximate response curves of datasets identified as significant by providing analytical results for the linearisation of a stable focus model, a simplification of the Wilson-Cowan model in the stable focus regime. Additionally, we fitted the full non-linear Wilson-Cowan model to these datasets, and we show that in each case the model can fit to the dynamics of patient tremor as well as to the phase response curve. The vast majority of top fits are stable foci. The model provides satisfactory prediction of how patient tremor will react to phase-locked stimulation by predicting patient amplitude response curves although they were not explicitly fitted. We report that the non-linear Wilson-Cowan model is able to describe response to stimulation more precisely than the linearisation.

## 1 Introduction

Essential tremor (ET) is the most common movement disorder, affecting 0.9% of the population [1]. It predominantly manifests as a tremor of the upper limbs, and can severely affect daily-life. When medications are ineffective or not tolerated, thalamic deep brain stimulation (DBS) is a well-established therapy option. Clinically available DBS continuously delivers high-frequency (≈ 130 Hz) electrical stimulation to deep structures within the brain via an electrode connected to a pulse generator implanted in the chest. There is no agreement in the research community on the mechanisms of action of high-frequency DBS [2], but it is believed there is room for improvement in terms of efficacy, decrease in power usage, avoidance of habituation, and most importantly reduction of side effects [3]. Reported side effects of high-frequency thalamic DBS include speech impairment (dysarthria), gait disorders, and abnormal dermal sensations (paresthesia) [4].

Because side-effects are the main clinical bottleneck, improving high-frequency DBS generally means stimulating less by closing the loop on a signal related to motor symptoms, while maintaining clinical efficacy. One example of closed-loop DBS is adaptive DBS, whereby stimulation is triggered in Parkinson’s disease (PD) patients when pathological neural oscillation amplitude in the beta band is higher than a threshold. Compared to high-frequency DBS, it has been shown to improve motor performance, and reduce speech side-effects in humans [5, 6, 7]. Another example is phase dependent stimulation, which has been investigated in a computational model of PD [8], and in PD patients [9, 10]. Phase-locked DBS has recently been studied as a new therapy for ET [11]. Hand tremor is recorded, and the reduction in stimulation comes from stimulating with a burst of pulses according to the phase of tremor, only once per period of the tremor rather than continuously. In some patients, the strategy only requires half the energy delivered by high-frequency DBS for the same effect. Optimising phase-locked DBS requires a detailed understanding of the phase dependence of the response across patients, but so far data collection from phase-locked stimulation experiments has been restricted to small datasets because patients fatigue quickly. While direct analysis of the data has proven insightful [11], modelling phase-locked stimulation would allow predictions to be made from analytic and computational studies regarding the phase dependence of the response to stimulation, and would open the door to supplement scarcely available patient data with synthetic data. The ability to easily generate large amounts of synthetic data could come in handy to help devise and test control algorithms, or when trying to predict an effect that, because of noise in recordings, can only be deciphered when a large number of trials is available.

Tremulous hand movements are believed to be closely related to thalamic activity [12, 13], and it is believed that ET originates in the cerebellar-thalamic-cortical pathway [14]. However detailed knowledge of how ET comes about is missing, which makes simple, canonical models natural candidates to study ET. Recently, phase-locked DBS was studied using Kuramoto phase oscillators which do not model interacting neural populations with distinct properties [15]. In the present work, we focus on a neural mass model, the Wilson-Cowan (WC) model, whose architecture can be mapped onto neural populations thought to be involved in the generation of ET, and allows for strong coupling between the populations. Additionally, stimulation can be delivered in the model to the most common stimulation site for ET, the ventral intermediate nucleus (VIM). The model describes the firing rates of an excitatory and an inhibitory population, and only has a few parameters, which makes it less prone to overfitting and significantly easier to constrain than more detailed models. The WC model has been shown to be adept at describing beta oscillations in PD [16, 17]. Moreover, the work presented in [18] provides evidence that the effects of high-frequency DBS for ET in a WC model are similar to the description given by conductance-based models. While the WC model has been used to design closed-loop strategies for PD [19, 20], whether a firing-rate model such as the WC can model the effects of phase-locked DBS has not been approached in the literature. Based on strong assumptions, Polina et al. reduced a WC model to a one-dimensional ordinary differential equation and looked at periodic forcing, but not in the context of DBS, and without attending to dependence on the phase of stimulation [21]. The present work will focus on reproducing the phase-dependent effects of phase-locked DBS measured in human data with a WC model.

Stimulation changes the phase and the amplitude of tremor and the dependence of these changes on the phase of stimulation can be quantified by the phase response curve (PRC, in this study change in tremor phase as a function of tremor phase) and the amplitude response curve (ARC, in this study change in tremor amplitude as a function of tremor phase). The ARC directly measures the change in tremor, hence the change in patient handicap, but both the ARC and the PRC are important to understand the effects of phase-locked DBS and potentially optimise the stimulation pattern. Theoretically, PRCs and ARCs have been defined differently, mostly in the context of limit cycle models concerned with asymptotic response to infinitesimal perturbations, see for example [22, 23, 24, 25, 26]. In patients, DBS stimulation is not infinitesimal, and tremor data is very variable so stimulation happens in transient states. Therefore rather than considering an asymptotic description of the changes in phase and amplitude, we will be focusing on a close variant of the experimental response curve measurement methodology applied to blocks of stimulation in [11], which we will hereafter refer to as the “block method”. It provides a finite time response to a finite perturbation and relies on the changes in the Hilbert phase and amplitude following blocks of phase-locked stimulation (more details in 2.1). The only exception to this will be in analytical derivations (section 4), where a first order measurement of the response curves (i.e. measurement at the end of the stimulation period) will be used for tractability, as a simplified first approach to the model. For coherence with the experimental response curve measurement methodology, the notion of phase and amplitude used throughout will be the Hilbert phase and amplitude or equivalent. It should also be noted that we are considering population response curves and not single neuron response curves. The vast majority of best performing WC models in reproducing patient data are found in this work to be stable foci, where tremor dynamics are being reproduced by adding noise to the system, so we restrict our analytical considerations to stable foci.

The main contributions of this work are the following. We first analyse patient response curves, identify a subset of datasets passing appropriate statistical tests, and characterise the relationship between PRC and ARC in these patients (section 2). Following the introduction of our biologically motivated WC model (section 3), we derive approximate analytical expressions that delineate the response to stimulation of a 2D dynamical system described by a linearised focus, with the goal of better understanding the constraints built in the model (section 4). The derived response curves are close to sinusoidal, and a relationship between them is found, revealing similarities in shape and phase shift with patients who have significant PRCs and ARCs. We then show that for these patients, the WC model can be fitted to the data and can reproduce the dependence of the effects of stimulation on the phase of stimulation. The model is fitted to the PRC and can reasonably predict the ARC, and notably what is approximately the best phase to stimulate (section 5). We then proceed to compare the relationship between response curves in the linearised and the full model and conclude that non-linearity is important to better reproduce the relationship found in patients (section 6). Finally a discussion is provided (section 7).

## 2 Patient response curves and their relationship

In order to assess phase dependence of the effects of DBS in patients, we extract PRCs and ARCs from patient’s tremor data, provide a statistical analysis of the curves, and analyse their relationship when applicable.

### 2.1 Analysis method

In the study reported in [11], ET patients are fitted with an accelerometer to record their tremor, and DBS locked to the phase of tremor acceleration is provided in blocks of 5 s to the VIM of the thalamus, with 1 s without stimulation between blocks. Each block targets a stimulation phase randomly selected out of 12 tremor phases (e.g. 120 degrees for the block shown in Figure 1). Stimulation is delivered once per period at the target phase, in the form of a burst of 4 to 6 pulses at high frequency (130 Hz or higher). There are about 10 trials available per phase (about 120 blocks per patient). The method described in [11] to obtain patient’s response curves was specifically developed for this type of data, and we closely follow it. We refer to our version of the method as the “block method” and denote the response curves obtained by bPRC and bARC, “b” standing for block. More specifically, we define the bPRC and the bARC according to the following procedure. Tremor frequency is around 5 Hz, and the dominant axis tremor recordings are bandpass-filtered (4 Hz band encompassing the patient tremor frequency content) by means of a backwards and forwards Butterworth second order filter (zero-phase filtering) and z-scored.

**Figure 1:**
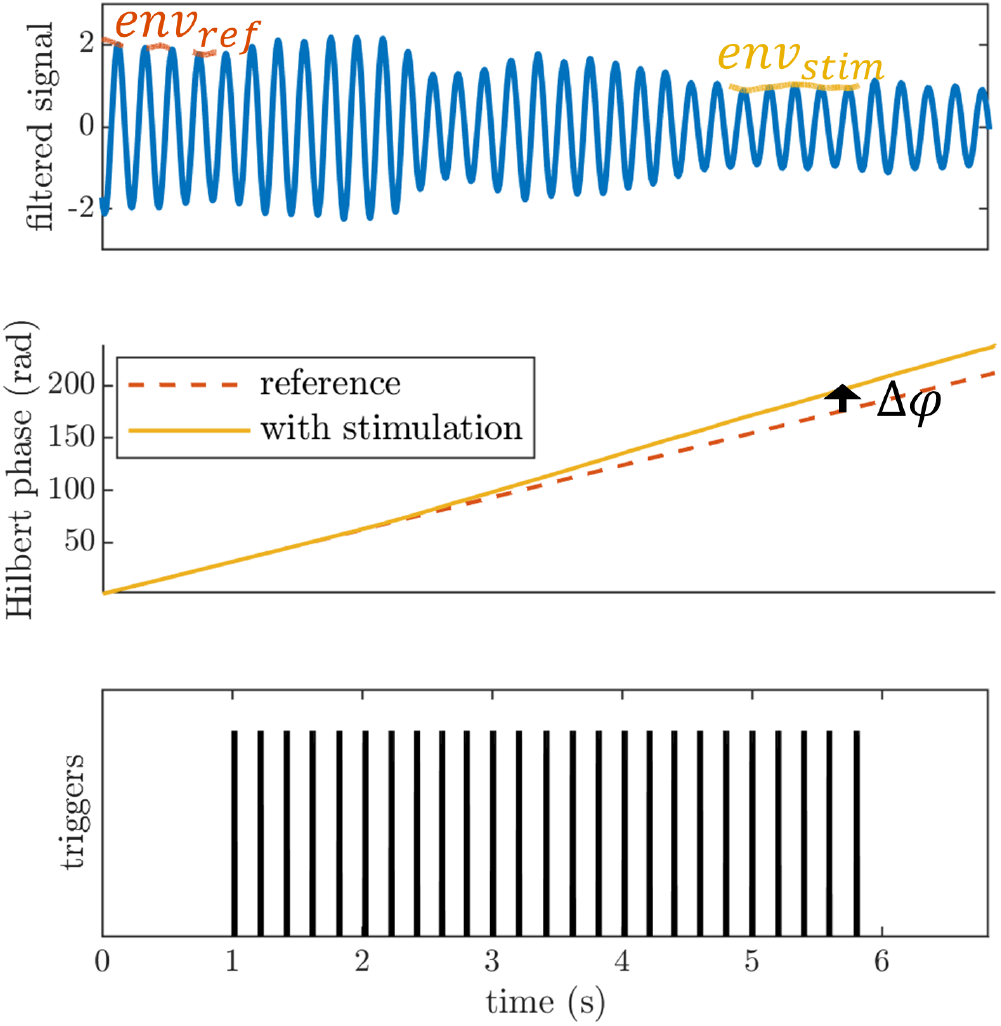
Example showing the block method applied to a block of stimulation with a target stimulation phase of 120 degrees. Stimulation triggers are shown in black in the lower panel while filtered tremor is shown in blue in the upper panel. The change in phase due to stimulation Δ*φ* is obtained by comparing at the end of the block the actual Hilbert phase to a linear phase obtained by a straight line fit to the phase evolution 1 s before the block (middle panel). The change in amplitude is given by the difference between the mean of *env_stim_* and *env_ref_* (top panel). Both the phase and amplitude responses are normalised by the number of pulses in the block.

#### Obtaining the change in phase (bPRC)

For each block, a straight line is fitted to the evolution of the Hilbert phase during the 1 s period without stimulation before the block. The change in phase Δ*φ* due to the block is given by the difference between the phase of the fitted reference line evaluated at the end of the block and the actual Hilbert phase at the end of the block (see Figure 1). This phase response is divided by the number of pulses in the block (on the basis of 4 pulses per burst for patient 4R and 4L, and 6 pulses per burst for the rest), which gives an average response for one pulse. The target phase at which stimulation is supposed to occur is known for each block, but phase tracking not being perfect, the actual Hilbert phase at which stimulation occurred is determined for each burst of stimulation as the circular mean of the Hilbert phase during the burst. We take the circular mean of these burst angles for a given block as the actual mean phase of stimulation for the block. These values are then binned into 12 phases bins, and the change in phase is averaged within bins to obtain the bPRC.

#### Obtaining the change in amplitude (bARC)

For each block, the change in amplitude is given by the difference between the mean of the Hilbert amplitude during the last second of the block and the mean of the Hilbert amplitude during the 1 second without stimulation before the block (see Figure 1). As for the change in phase, this amplitude response is divided by the number of pulses in the block, and averaged across blocks in the same phase bin to obtain the bARC.

#### Measuring response curves significance and PRC-ARC shift

In order to identify significant patient’s response curves, we performed two statistical analysis. First, bPRCs and bARCs were tested for a main effect of phase by means of a Kruskal-Wallis ANOVA (12 phase bins) to differentiate patients’ response curves that may be dominated by noise (lack of phase-dependent response or data collection and analysis unable to measure it). Second, since we are expecting response curves to have a dominant first harmonic, the cosine model *y* = *c*_1_ + |*c*_2_| cos(*x* + *c*_3_) was fitted to patients’ phase response and amplitude response curves. We assessed via F-tests whether the cosine model was better at describing the data than a straight line at the mean. Including the less specific ANOVA test allows for more generality, as we do not wish to exclude patients with significant, but non-sinusoidal response curves. On the other hand, the cosine test is more likely to detect phase-dependent effects of stimulation in patients which indeed have sinusoidal response curves. We therefore define the following criterion for selection of a patient for further study in the rest of the manuscript.

#### Significance criterion

*having both bPRC and bARC deemend significant under FDR control (see below) by at least one of the two tests – ANOVA test for a main effect of phase or cosine model F-test*.

In both cases, the adaptive linear step-up procedure modified by Storey et al. in [27] and reviewed in [28] was used to keep the false discovery rate (FDR) below 5%. It improves on the original Benjamini and Hochberg procedure [29] by using an estimator 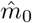 for the number of true null hypothesis *m*_0_ (total number of tests *m* = 2 response curves × 6 patients = 12 for each analysis). Controlling the FDR at 5% guarantees that the expectation of the number of false positives over the number of positives is less than 5%.

Additionally, in datasets where both bPRC and bARC are significant according to the cosine F-test, the relationship between bPRC and bARC is quantified by the shift in phase between the cosine model fits to the bPRC and the bARC. In these datasets, the PRC-ARC shift is calculated as 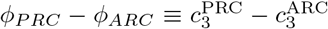 (mod 2*π*), with *ϕ_PRC_* – *ϕ_ARC_* ∈ [0, 2*π*). Calculating a PRC-ARC shift in other cases is not meaningful.

### 2.2 Results of the analysis

We analysed six datasets from the five patients included in [30] (datasets 4R and 4L are for the right and left upper limbs of the same patient). bPRCs and bARCs obtained are shown in supplementary Figure 15 in Appendix H, and results of the statistical tests are presented in Table1. Based on the significance criterion defined in the previous section, patients 1, 5 and 6 are selected for further study, as both their bPRCs and bARCs are found significant by the cosine F-test under FDR control. We note that patient 5 also has both his response curves deemed significant by the ANOVA test under FDR control. Datasets 3, 4R and 4L do not satisfy our selection criterion. In Figure 2, the shift *ϕ_PRC_* – *ϕ_ARC_* is plotted for patients for whom the cosine model was deemed significant in describing both their bPRC and bARC (which happens to be the same subset as patients satisfying our significance criterion). They have a shift in 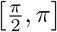, patients 5 and 6 being quite close to 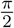.

**Table 1:**
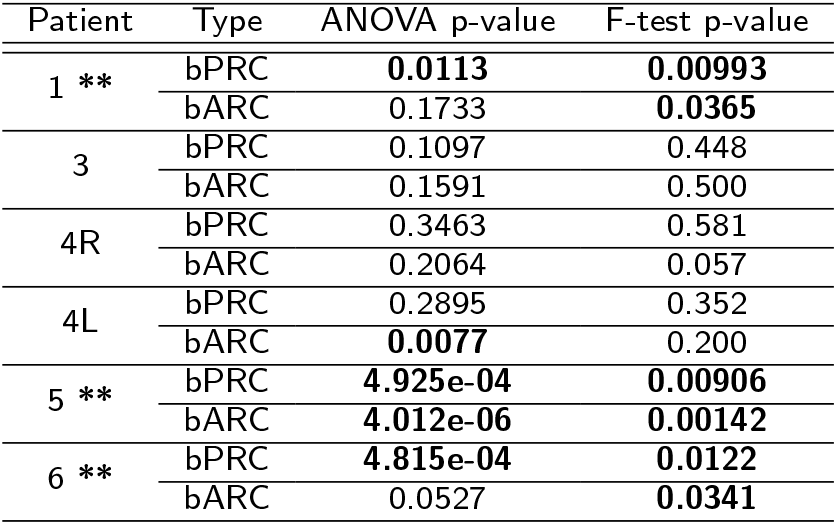
P-values of both statistical tests performed on patients’ response curves: Kruskal-Wallis ANOVAs testing a main effect for phase in patients’ response curves (third column), and cosine model F-tests (fourth column). P-values in bold are deemed significant with FDR control at the 5% level (separate FDR analyses per test type, 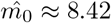 for the ANOVAs and 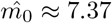 7.37 for the F-tests). Double stars indicate datasets satisfying our significance criterion as defined in section 2.1.

| Patient | Type | ANOVA p-value | F-test p-value |
| --- | --- | --- | --- |
| 1 ** | bPRC | **0.0113** | **0.00993** |
|  | bARC | 0.1733 | **0.0365** |
| 3 | bPRC | 0.1097 | 0.448 |
|  | bARC | 0.1591 | 0.500 |
| 4R | bPRC | 0.3463 | 0.581 |
|  | bARC | 0.2064 | 0.057 |
| 4L | bPRC | 0.2895 | 0.352 |
|  | bARC | **0.0077** | 0.200 |
| 5 ** | bPRC | **4.925e-04** | **0.00906** |
|  | bARC | **4.012e-06** | **0.00142** |
| 6 ** | bPRC | **4.815e-04** | **0.0122** |
|  | bARC | 0.0527 | **0.0341** |

**Figure 2:**
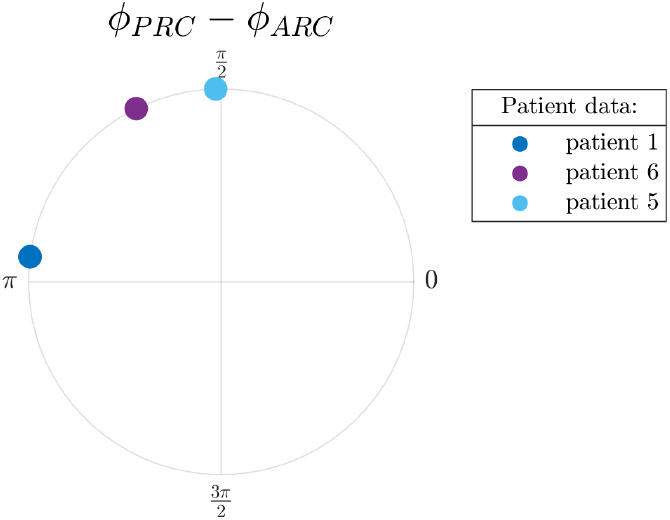
PRC-ARC shift in patients. Only showing patients with significant cosine model F-test for bPRC and bARC under FDR control. The calculated PRC-ARC shifts are in 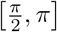.

## 3 Implementation of the Wilson Cowan model for essential tremor DBS

To model the experimental data described in the previous section, we use a WC model that describes the interaction between an excitatory and an inhibitory population of neurons. Specifically, we map a 2 population WC model without delays as described in [31] onto the anatomy of the thalamus (Figure 3). The circuit we are about to describe is a good candidate, but not the only biologically plausible mapping of an excitatory/inhibitory loop in the context of tremor. In our candidate mapping, the VIM is modelled as an excitatory population, connected to an inhibitory population of the thalamus, the reticular nucleus (nRT). The high coherence between ventral thalamic activity and electromyographic recording of the contralateral wrist flexors [12, 13] is our justification for modelling tremor by the activity of the excitatory population. VIM and nRT are reciprocally connected (the excitatory projections from VIM to nRT are via Cortex). The VIM receive a constant input from the deep cerebellar nuclei (DCN) and is part of a self-excitatory loop via Cortex. nRT receives a constant cortical input. We add Gaussian white noise to this two-population WC, and the activity of the VIM, *E*, and the activity of the nRT, *I*, are described by the stochastic differential equations

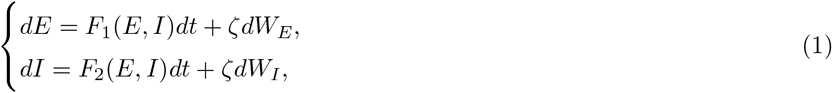

where *dW_E_* and *dW_I_* are Wiener processes, *ζ* the noise standard deviation. We define

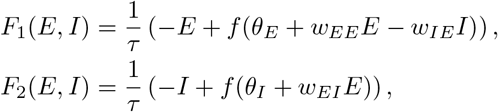

with *w_PR_* the weight of the projection from population “P” to population “R”, *θ_P_* the constant input to population “P”, *τ* a time constant (assumed to be the same for both populations). We use a sigmoid function

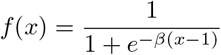

parametrized by a steepness parameter *β* (same choice as in [31]). The VIM is the most common target of DBS for ET, which is why we model stimulation as a direct increase in *E*. Analytical expressions for response curves are out of reach for the full non-linear model, and we will study next a linearisation of a deterministic stable focus model.

**Figure 3:**
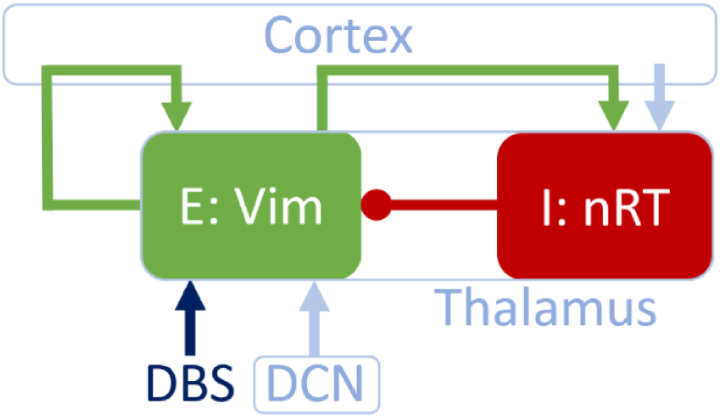
The WC model can describe the populations thought to be involved in the generation of ET. The excitatory population E and the inhibitory population I model respectively the VIM and the nRT of the thalamus. Arrows denote excitatory connections or inputs, whereas circles denote inhibitory connections. The VIM is the target of DBS and also receives an input from the deep cerebellar nuclei (DCN). The self-excitatory loop of the VIM, as well as the excitatory connection from VIM to nRT are via cortex.

## 4 Response curves and their relationship in a focus model

The goal here is to derive approximate analytic expressions for the first order phase and amplitude responses to one pulse of stimulation in a 2D dynamical system that is described by a (stable) focus. Such a linearisation can be applied to the deterministic WC model given by equation (1) with *ζ* = 0 in the focus regime. We follow the previous section in modelling the tremor signal as the first coordinate of the dynamical system, and in providing stimulation pulses along the first dimension. The results will provide a basis for understanding how the effects of stimulation on phase and amplitude are coupled in the WC model, and for comparison with experimental data.

### 4.1 Linearisation of a focus

To distinguish scalars and vectors more easily, vectors will be denoted in bold. Let **Ž** = *F*(**Z**) be a dynamical system, where **Z** ∈ ℝ^2^ and *F* is differentiable. The Jacobian of *F* is

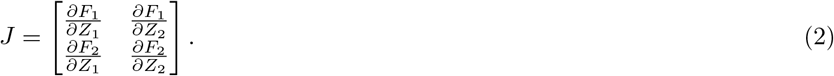

Let **Z\*** be a fixed point of *F*. The dynamics of **X** = **Z** − **Z\*** are approximated in the vicinity of the equilibrium X = 0 by the linear equation

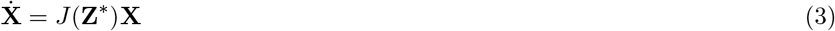

where *J*(**Z\***) is the Jacobian evaluated at the fixed point. We will treat the case of Jacobians having complex conjugate eigenvalues λ_±_=*σ*±*iω*. In particular, we are interested in stable foci, which imply *σ* < 0. The WC model can operate in that regime [31]. The case of centers (*σ* = 0, purely imaginary complex conjugate eigenvalues) will also be described, although it is of little interest for patient fits. If **k** = **a** + *i***b** is the right eigenvector associated with λ_+_, *K* and *K*′ coefficients determined according to initial conditions, the general real valued solution of (3) reads

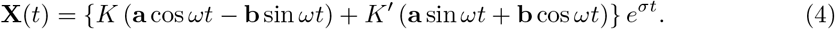

We will be using the following notations for the coordinates of the eigenvector:

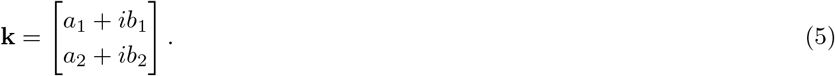

Equation (4) and what follows are not valid in the case of multiple or repeated real eigenvalues, which are of no interest for our purposes (no rotation).

### 4.2 Phase definition

The notion of phase is central to phase-locked stimulation, and in this section we define phase in a way that is approximately equivalent to what is commonly used in the analysis of experimental data. A typical signal only has one component, and the Hilbert transform provides a convenient way of reconstructing a phase from a single component. Despite being a protophase (see discussion section), the Hilbert phase is widely used to analyse experimental data (see for instance [9, 11, 32, 33, 34]), and this is the reason why we choose in our linearised system a phase definition approximately equivalent to it. We define a phase variable as *ϕ*= *ωt* with a zero phase point defined as the maximum of *X*_1_(*t*) (similarly to the Hilbert phase), which is therefore on the nullcline of the first coordinate. This phase definition is different from other common definitions such as the trajectory polar angle in the phase plane of a 2D system, or isochronal (asymptotic) phase. We demonstrate next that it is very close to the Hilbert phase of *X*_1_ for slow decay. It should be noted that this is generally only true for the linearisation. As the Hilbert phase is also the phase definition used in the other sections of this manuscript, the following proof ensures consistency.

Let 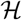 denote the Hilbert transform. To establish equivalence of our phase definition with the Hilbert phase of *X*_1_ given by

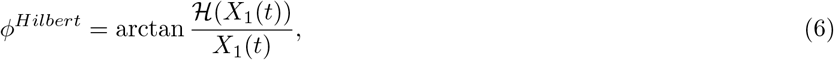

a first step is to calculate the Hilbert transform of the signal *X*_1_(*t*). The Hilbert transform is a linear operator, and *X*_1_(*t*) is a linear combination of *s*(*t*)*s_c_*(*t*) and *s*(*t*)*s_n_*(*t*) with *s*(*t*) = *e*^*σ*|*t*|^, *s_c_*(*t*)= cos *ωt*, and *s_n_*(*t*)= sin*wt* (see equation (4)). Inspired by the proof of the Bedrosian identity [35], we calculate error terms and show in Appendix A that 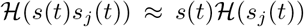 for *j* = *c,n*. The Hilbert phase of *X*_1_ is therefore given by

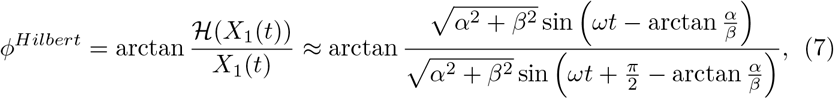

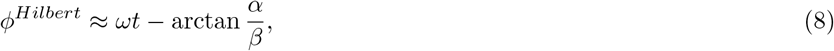

where

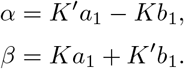

In the case of *ω* ≫ |*σ*|, it can be shown that *α* and *β* are such that *ϕ^Hilbert^* is referenced to the maximum of *X*_1_.

### 4.3 Reference trajectory and stimulated trajectory

In order to calculate first order response curves, we will consider a reference trajectory without stimulation, and a trajectory that underwent an instantaneous stimulation pulse *δX*_1_ at a stimulation phase *ϕ*_0_. The effects of stimulation on phase and amplitude will be measured at the next maximum of *X*_1_ for both trajectories. We will denote these hPRC^(1)^ and hARC^(1)^ as they are first order responses based on a phase definition equivalent to the Hilbert phase. A sketch of the method is provided in Figure 4.

Expressions for the coefficients *K_ref_* and 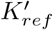 of the reference trajectory are derived in Appendix B. We want to study the effects of stimulating at phase *ϕ*_0_. The point of stimulation **X**^1−^ at phase *ϕ*_0_ is expressed as

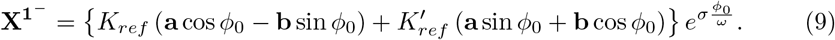

An instantaneous stimulation *δX*_1_ is applied at **X**^1−^ as

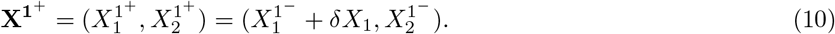

The trajectory after stimulation is still constrained by the dynamics given by equation (4), which allows for expressions for the coefficients on this new trajectory *K_stim_* and 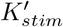 to be found (see Appendix C). To measure the change in phase and amplitude between the next peaks of the stimulated trajectory and the reference trajectory, the phase *ϕ_max_* of the next maximum of the first coordinate on the stimulated trajectory 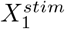 is needed (the phase of the next maximum of *X*_1_ on the reference trajectory is 2*π*). A derivation for *ϕ_max_* is provided in Appendix D.

### 4.4 Phase response

The first order phase response curve can be calculated based on the reference trajectory period *T*_0_ and the stimulated trajectory period *T_stim_*, which is given by the sum of the time spent on the reference trajectory before stimulation and the time spent on the new trajectory after stimulation:

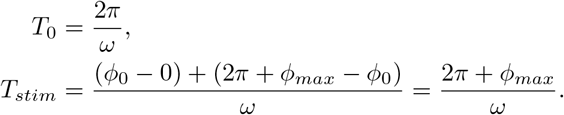

**Figure 4:**
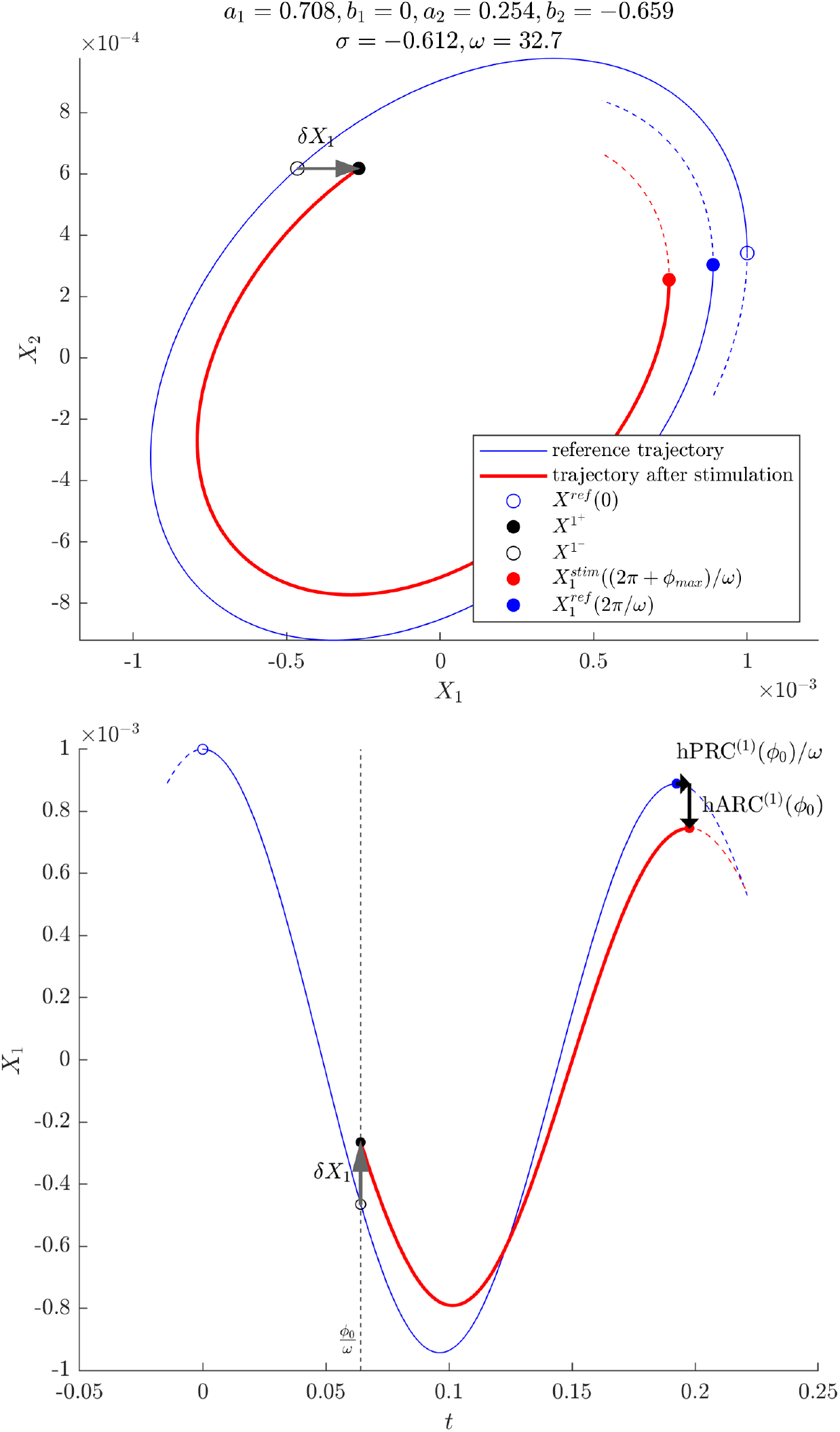
Illustration of the approach taken to derive expressions for the phase and amplitude responses in the linearisation of a 2D focus model. Top: phase plane, bottom: time-series of *X*_1_. The tremor is modelled by *X*_1_, and the stimulation *X*_1_ is applied to *X*_1_. The system shown corresponds to the linearised fit of patient 1 as described in section 6.1.

For a phase response curve in radian, we obtain

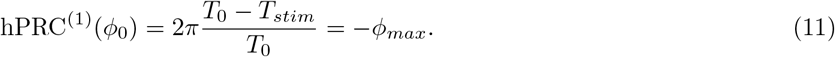

The phase *ϕ_max_* depends on *δX*_1_ through *K_stim_* and 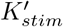, and a Taylor expansion around 0 yields, to lowest order in *δX*_1_,

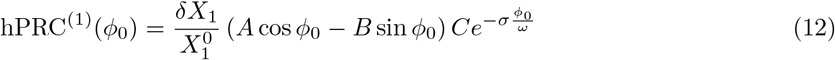

with

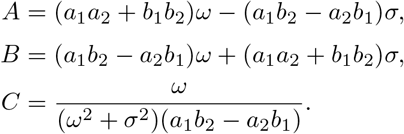

Although we are omitting the amplitude dependence in our notations for convenience, the first order PRC is found to be proportional to the inverse of the peak amplitude of the oscillations at the beginning of the stimulation period 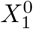. It is also directly proportional to the stimulation amplitude *δX*_1_, and directly depends on phase via sinusoidal functions and a factor related to the decay. The constants *A, B*, and *C* only depend on the real and imaginary parts of the eigenvalue λ_+_ and the associated eigenvector **k**.

### 4.5 Amplitude response

For our purposes we are interested in the amplitude of the first coordinate, and the first order amplitude response curve (ARC) is obtained as the difference in first coordinates between the stimulated and the reference trajectories evaluated at their respective next peak after stimulation. It should be noted this is equivalent to a first order change in Hilbert amplitude, at least for *w* ≫ |*σ*|. The first order ARC is calculated as

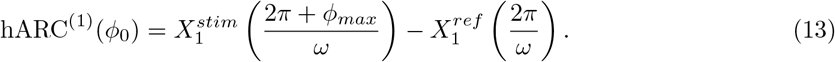

A Taylor expansion around 0 yields, to lowest order in *δX*_1_,

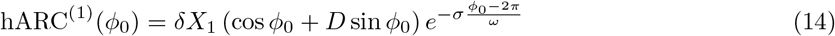

with

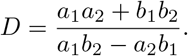

Interestingly, the first order ARC close to the fixed point does not depends on the amplitude of the oscillations 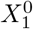. As expected, the first order ARC is directly proportional to the stimulation amplitude *δX*_1_. Similarly to the first order PRC, it directly depends on phase via sinusoidal functions and a factor related to the decay, and the constant *D* only depends on **k**. The obvious similarities between the first order PRC and ARC suggest there may be a relationship between the two.

### 4.6 Relationship between first order PRC and ARC

We seek a relationship involving the derivative of the first order PRC, which, based on equation (12), is given by

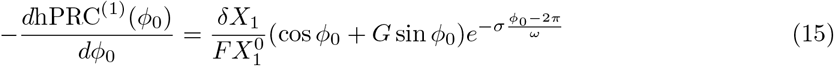

with

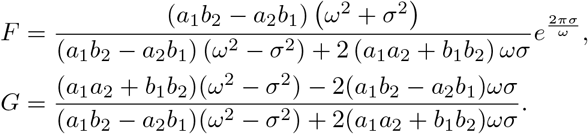

Let us study the case where *ω* ≫ |*σ*|:

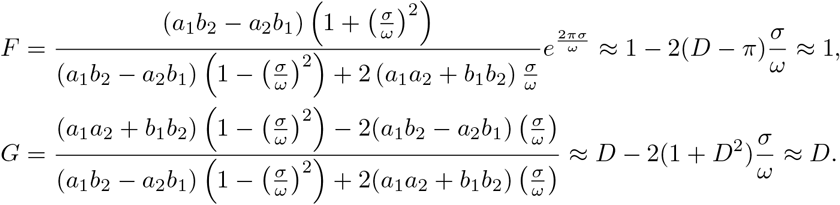

Therefore in that case the first order ARC is approximately the opposite of the derivative of the first order PRC scaled by the peak amplitude at the beginning of the stimulation period (in general, the scaling factor is 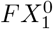):

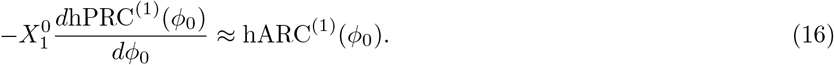

For a slow decay compared to the rotation, the PRC-ARC shift in the linearisation of a focus will therefore be close to 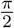, which is the value observed for patient 5 (see Figure 2). A detailed analysis of the PRC-ARC shift in the model is provided in section 6.

### 4.7 Applications to simple systems

We turn to simple systems to illustrate the results of the previous sections. In what follows, response curves are given for *δX*_1_ =2 × 10^−4^ and 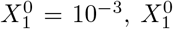 being a maximum of *X*_1_ as a function of time (these only act as scaling factors of the response curves and will not change their shape).

#### Circular flow without decay

As an introductory example, let us consider a simple circular flow for which the *J* matrix is

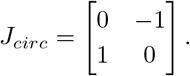

The eigenvalues of *J_circ_* are ±*i* so the results of the previous sections can be applied. Equations (12) and (14) are plotted for this system with our choice of *δX*_1_ and 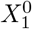. The result for the first order PRC is shown in Figure 5, panel A2, and for the first order ARC in panel A3. For this system, hPRC^(1)^ is simply the opposite of a sine, hARC^(1)^ simply a cosine. Moreover, *σ* = 0, hence *G* = *D* (see section4.6) and equation (16) is exact, as exemplified in Figure 5, panel A3. hARC^(1)^ is obtained by only scaling the derivative of hPRC^(1)^ by 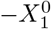 as *a*_2_=*b*_1_=0 and *F*=1. Note that WC parameters for which the system’s Jacobian at the fixed point is *J_circ_* cannot be found as the second diagonal term cannot be 0, at least in the version of the WC model used in this work (see (38) in Appendix E).

**Figure 5:**
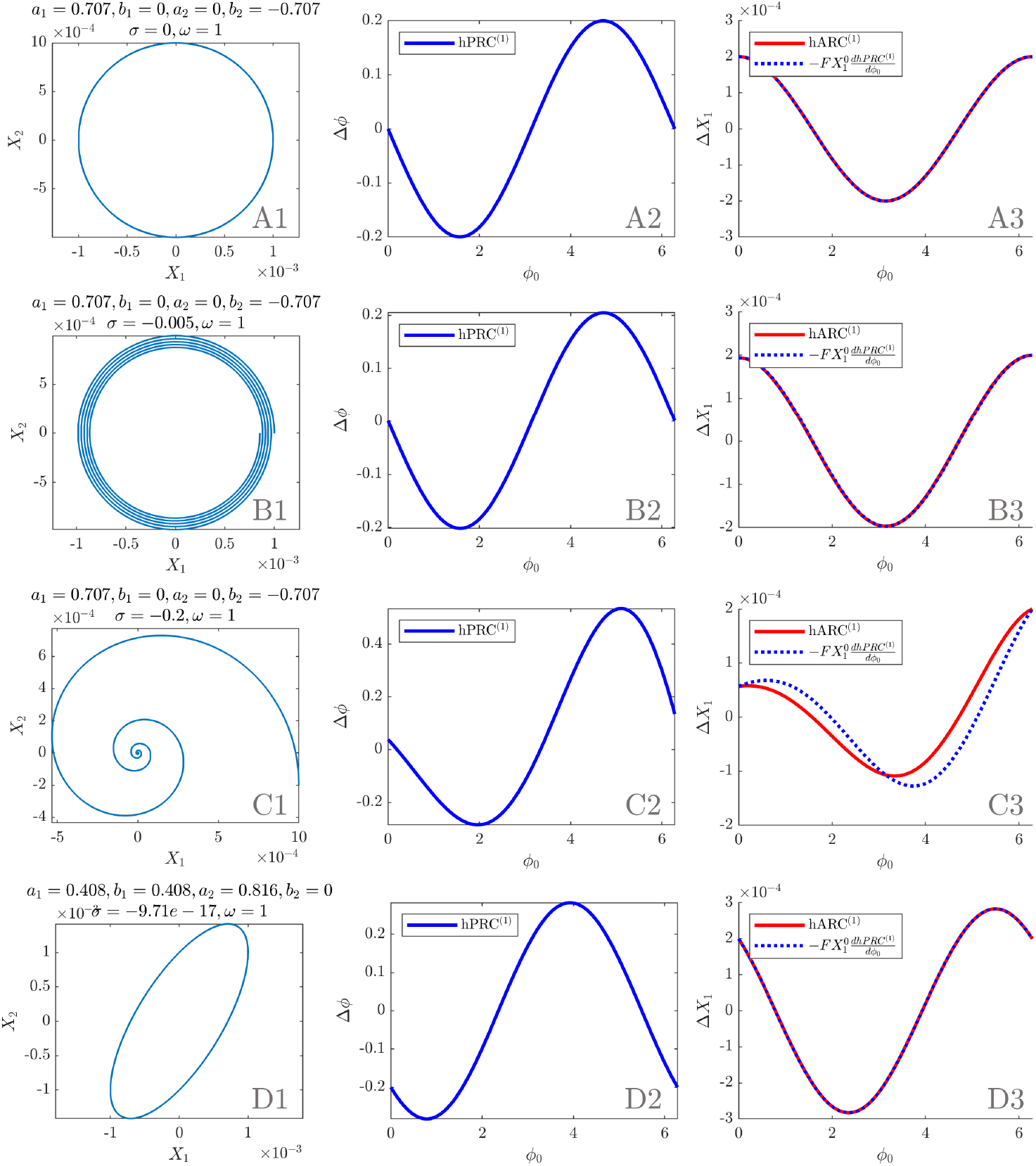
Analytical results in simple systems (initial conditions as in the main text). First column: phase space. Second column: first order PRC as per equation (12). Third column: first order ARC as per equation (14) and opposite of the derivative of the first order PRC scaled by 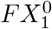. Panel A corresponds to *J_circ_* (circular flow, no decay), panel B to 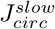 (circular flow, slow decay), panel C to 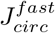 (circular flow, fast decay), and panel D to *J_ellip_* (tilted elliptic flow, no decay).

#### Circular flow with decay

We can introduce a slow decay (Figure 5, panel B) and then a fast decay (Figure 5, panel C) in the circular flow. We choose the *J* matrices

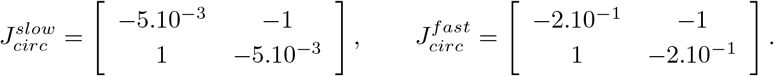

The slow decay leads to a scaling factor *F* ≈ 1, and the approximation of equation (16) is very good, as *ω* ≫ |*σ*| (see Figure 5, panel B3). The case of the fast decay corresponds to *ω* = 5|*σ*|. The first order PRC and ARC no longer look like pure sinusoids and the approximation relating the response curves is less accurate (*ω* = 200|*σ*|, see Figure 5, panel C3). It is possible to find WC parameters for which the system’s Jacobian at the fixed point is 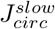 or 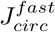. How such parameters are found is explained in Appendix E, and the results are presented in supplementary Table (3) in Appendix I. In both cases, *w_IE_* = *w_IE_*, and *w_EE_* = 0.

#### Tilted elliptic flow without decay

The tilted elliptic flow without decay of Figure 5, panel D, corresponds to the *J* matrix

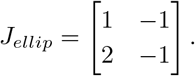

The first order PRC and ARC are sums of a sine and a cosine, which brings a horizontal shift in phase compared to a circular flow without decay. The eigenvalues are still purely imaginary, but *F* is no longer one. Because *σ* = 0, the relationship of equation (16) is still exact (see Figure 5, panel D3). It is possible to find WC parameters for which the system’s Jacobian at the fixed point is *J_ellip_* (see supplementary Table (3) in Appendix I). Patient fits fall in the category of (potentially tilted) elliptic flows with decay, and will be dealt with in section 6.1. The linearised stable focus model exhibits close to sinusoidal response curves and a PRC-ARC shift close to 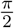, which when contrasted with patient data (response curves passing the cosine model F test and PRC-ARC shifts in 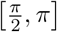 as shown in Figure 2), provides a strong motivation to fit the WC model to data.

## 5 Fitting the full Wilson Cowan model to patient data and response to phase-locked stimulation

### 5.1 Fitting procedure

We now turn to fitting our stochastic neural mass model (equation (1)) to patient data. The model is fitted to features (also known as summary statistics) extracted from patient tremor recordings. The parameters we fit are shown in Table 2, and include model parameters, stimulation magnitude and stimulation delay (time between the stimulation trigger is recorded and stimulation is actually provided to the E population, more about its interpretation in section 7). Stimulation is implemented directly in the Euler update of our integration scheme. We aim at reproducing tremor dynamics and fit to three dynamical features: the power spectrum density of the data (PSD), its Hilbert envelope probability density function (PDF), and its Hilbert envelope PSD. While the envelope PDF captures the range of amplitudes present in the tremor, the envelope PSD describes how quickly tremor amplitude changes. But we also aim at reproducing response to stimulation, and fit to the patient bPRC. The data dynamical features are obtained after filtering and z-scoring the data as described in section 2.1. The data bPRC is obtained as described in section 2.1.

**Table 2:** Best parameters for the 3 fitted patients.

| Parameter | Symbol | Best fit values |  |  |
| --- | --- | --- | --- | --- |
|  |  | Patient 1 | Patient 5 | Patient 6 |
| I to E weight | $w_{IE}$ | 9.4014 | 26.048 | 5.2064 |
| E to I weight | $w_{EI}$ | 9.6306 | 25.3384 | 24.4813 |
| E to E weight | $w_{EE}$ | 6.7541 | 1.548 | 2.7514 |
| Sigmoid steepness parameter | $\beta$ | 1.1853 | 2.4234 | 4.1933 |
| Time constant (s) | $\tau$ | 0.0758 | 0.29984 | 0.2513 |
| Constant input to E | $\theta_E$ | 1.4240 | 22.8621 | 2.9127 |
| Constant input to I | $\theta_I$ | -3.2345 | -9.9279 | -3.4008 |
| Noise standard deviation | $\zeta$ | 0.0457 | 0.013707 | 0.0263 |
| Stimulation magnitude | $\delta E$ | 0.001684 | 0.00598 | 0.001686 |
| Stimulation delay (ms) | $\Delta t_{stim}$ | 138.8366 | 444.1573 | 183.4711 |

The fitting procedure is summarized in Figure 6. Local optimisations are carried out using gradient free optimisation, specifically a direct search algorithm called the generalized pattern search algorithm (more details are given in Appendix F). In order to measure response to stimulation as in the data, each local optimisation step needs to simulate the model with phase-locked blocks of stimulation. This requires integrating the differential equations of the model while tracking the phase and providing stimulation at the right time, which is done by monitoring the zero-crossing phase alongside a Euler integration scheme. Appendix G details implementation of the simulator. The four features are computed on the model output at each optimisation step. The same method is used as for the data features, with three differences. The first is that for increased stability of the optimisation, the model bPRC is averaged over a much greater number of trials (600 trials), while the more robust dynamical features are obtained from nine trials only to reduce computation cost. The second is that the model output is not filtered to compute the dynamical features (only z-scored), as we want the model output to primarily generate the filtered tremor signal (a model generating mostly 1Hz oscillations but reproducing patient tremor when filtered at 5Hz would not be desirable). Computing the bPRC still requires filtering, as it relies on the Hilbert transform. The third difference is that the filtering window for the bPRC cannot be adjusted manually in optimisation steps, so a 4Hz band centered on the model PSD peak is used. As for the data bPRC and bARC, the actual Hilbert phase at which stimulation occurred is used to compute response curves via the re-binning process described in section 2.1, and the zero-crossing phase is only needed to enable phase-locked stimulation in the model. Phase-tracking performance is illustrated in supplementary Figure 16 in Appendix.

**Figure 6:**
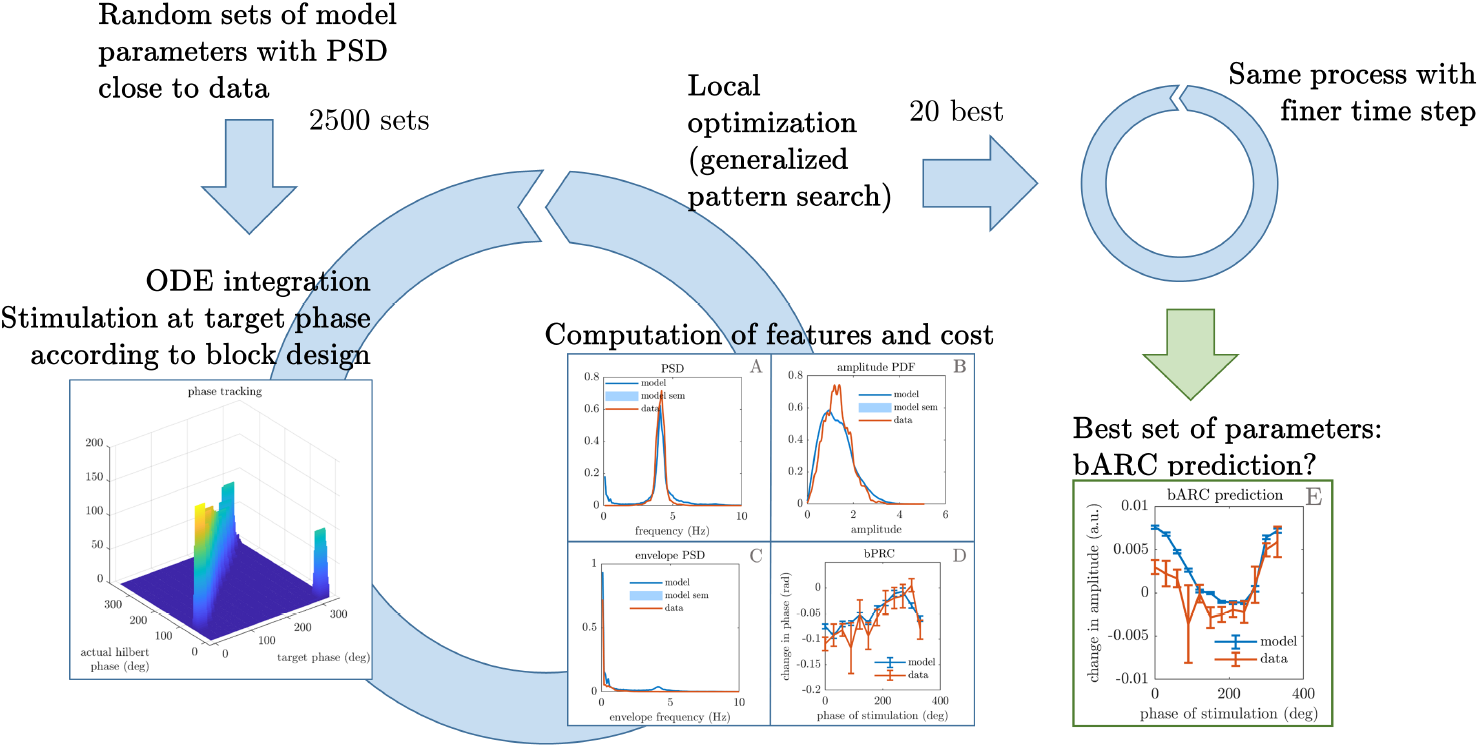
The fitting procedure involves 2500 local optimisations for each patient. The simulation of the model at each optimisation step requires to track the zero-crossing phase in order to provide stimulation at the right phase. The phase-tracking ability of the scheme is satisfactory when compared to the actual Hilbert phase (left, detailed in Figure 16 in Appendix H). The optimiser minimises a cost function that includes the comparison of three tremor dynamics features (tremor PSD, tremor envelope PSD, tremor envelope PDF) plus the PRC against the data (middle). Following a second optimisation of the 20 best results with a finer time step, a best set of parameter comes out of the procedure, and the model ARC can be compared against the data ARC. More details on the fitting procedure are given in Appendix F.

At each step, once the four features have been computed on the model output, the optimiser returns the cost

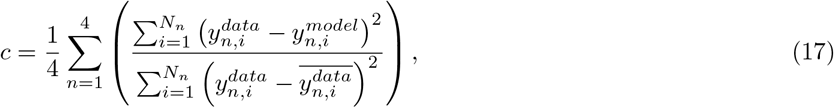

with *y_n_, n* ∈ {1, 2, 3,4} being the four features considered, and *N_n_* the length of *y_n_*. At the end of the procedure, the fit with the lowest *R*^2^ = 1 − *c* for each patient is deemed the best fit. In case of a tie (difference in mean costs lower than standard error of the mean), foci are preferred over limit cycles.

### 5.2 Results of the fits

Patients passing our significance criterion (section 2) are fitted to, namely patient 1, 5, and 6. For these patients, we find that the model successfully reproduces tremor dynamics, including tremors with sudden bursts, and can fit to patient phase response to stimulation. The best fits obtained upon completion of the optimisation procedure are shown in Figures 7, 8, and 9. In addition to reproducing tremor dynamics and being able to fit to patient bPRCs, the model seems to be able to reasonably predict patient bARCs (obtained as in section 2.1, but not fitted to), and in particular which phases are approximately the best phase to stimulate, i.e. the phases at which the maximum decrease in tremor happens. Because of averaging across 600 trials, the model bPRC and bARC error bars are small compared to the data error bars (only about 10 trials).

**Figure 7:**
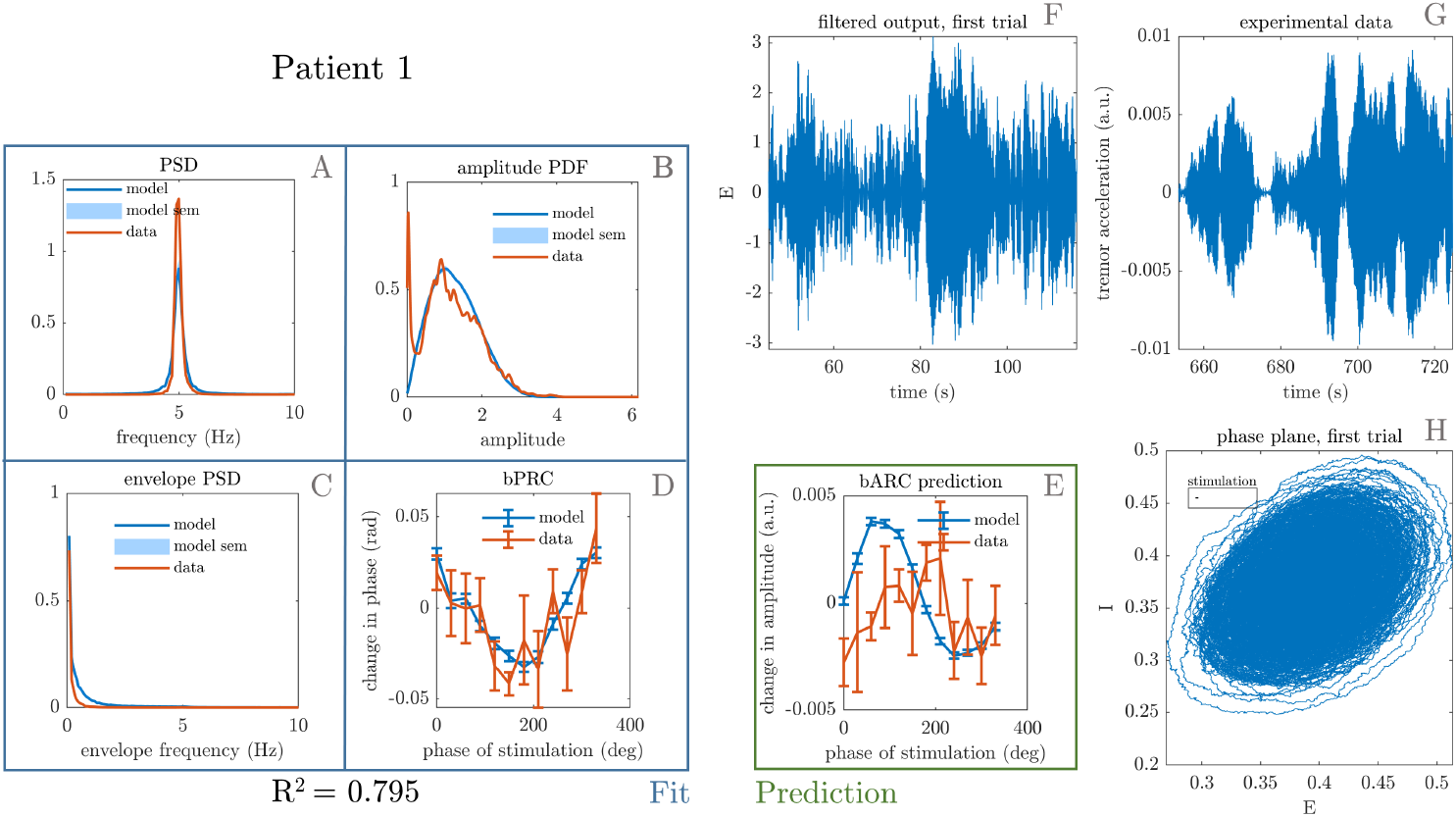
Best fit to patient 1. The four features that were included in the cost function are shown on the left, namely tremor PSD (A), tremor envelope PDF (B), tremor envelope PSD (C) and bPRC (D). The *R*^2^ for the model fit to these features is 0.795, and the model reasonably predicts the data bARC (E). The model phase plane is shown in H, and the model tremor time-series (F) is shown next to the patient tremor time series (G). The framed black bar in H indicates the fitted stimulation magnitude to scale.

**Figure 8:**
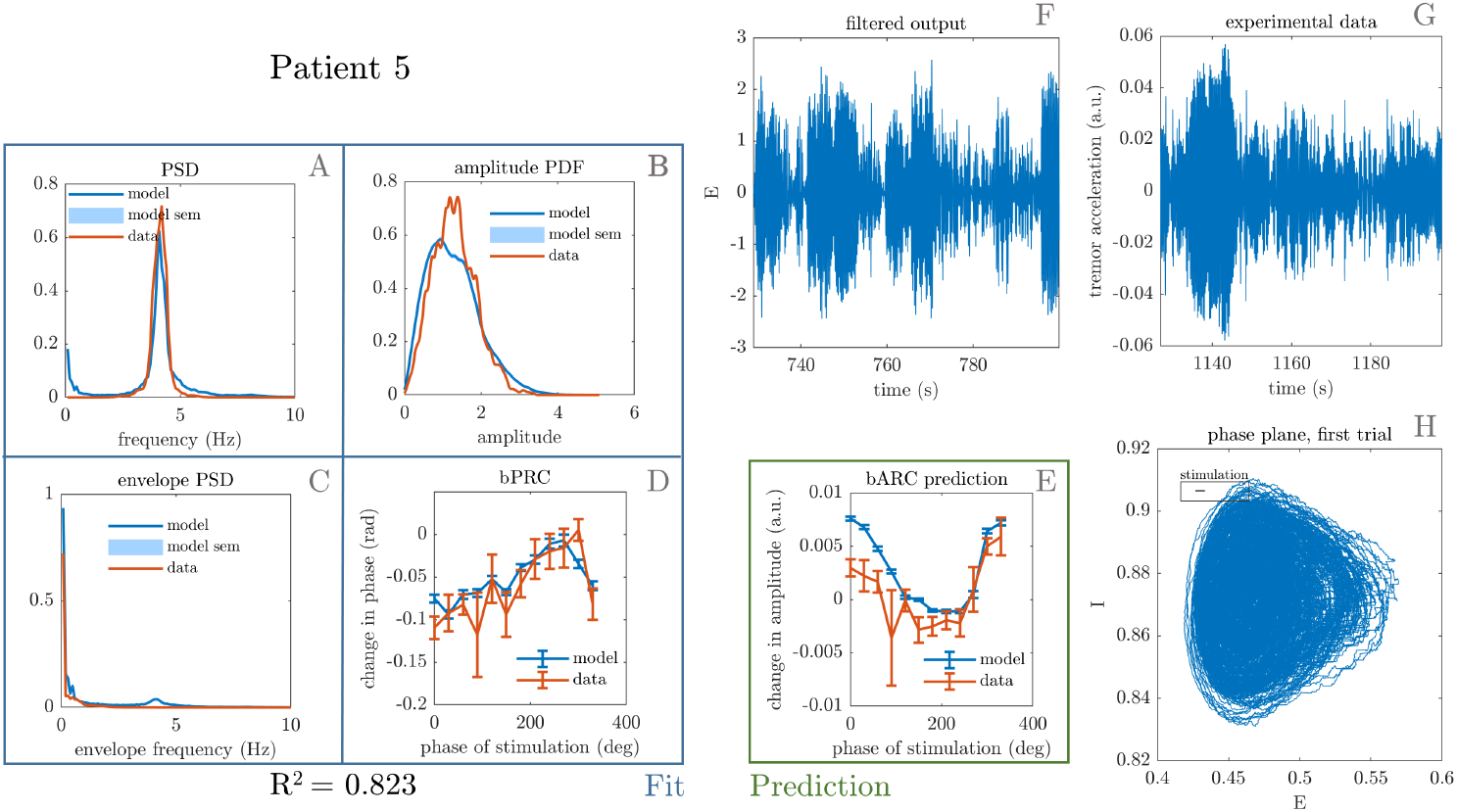
Best fit to patient 5. The four features that were included in the cost function are shown on the left, namely tremor PSD (A), tremor envelope PDF (B), tremor envelope PSD (C) and bPRC (D). The *R*^2^ for the model fit to these features is 0.823, and the model predicts the data bARC (E). The model phase plane is shown in H, and the model tremor time-series (F) is shown next to the patient tremor time series (G). The framed black bar in H indicates the fitted stimulation magnitude to scale.

**Figure 9:**
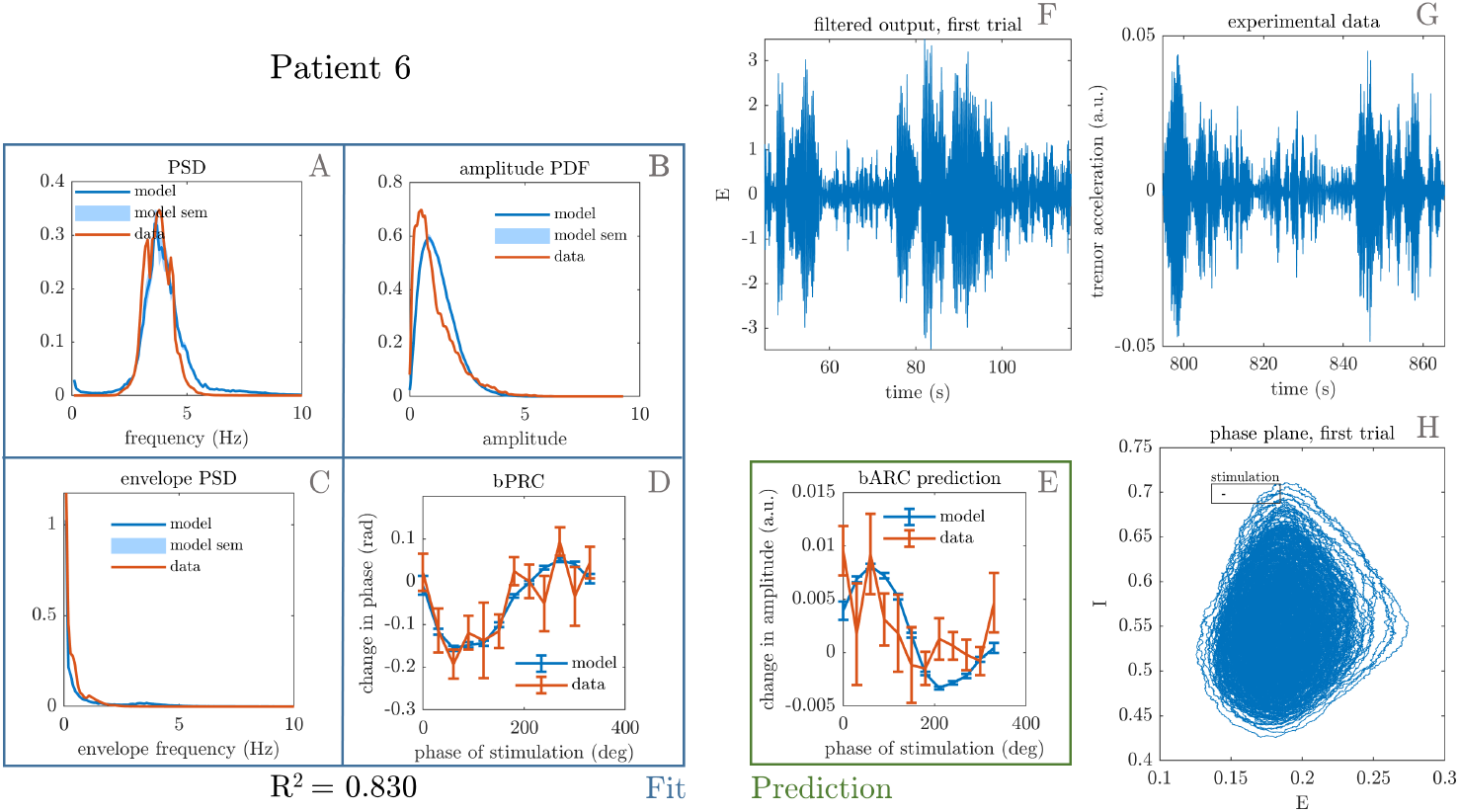
Best fit to patient 6. The four features that were included in the cost function are shown on the left, namely tremor PSD (A), tremor envelope PDF (B), tremor envelope PSD (C) and bPRC (D). The *R*^2^ for the model fit to these features is 0.830, and the model reasonably predicts the data bARC (E). The model phase plane is shown in H, and the model tremor time-series (F) is shown next to the patient tremor time series (G). The framed black bar in H indicates the fitted stimulation magnitude to scale.

#### Validating fitted stimulation magnitude

Cagnan et al. [11] report what the device settings are, and in particular the total electrical energy delivered (TEED) per unit time. We can build a quantity based on the fitted stimulation that should scale with the TEED per unit time. Because of z-scoring along the *E* dimension, we have to divide the fitted stimulation which is measured along the *E* dimension by the standard deviation of *E* (before z-scoring), and turn the fitted stimulation into an effective stimulation. As bursts are delivered once per period, this effective stimulation should be multiplied by the mean frequency of *E* to obtain a quantity proportional to energy per unit time (the number of pulses per burst is the same for the three patients). Figure 10 shows the effective stimulation times the mean frequency for the 15 best performing fits against the TEED per unit time for each patient (correlation coefficient for fit averages *r* = 0.98). Under the assumption that patient intrinsic sensitivities to stimulation are somewhat similar, we can conclude from the correlation that the fitting procedure successfully captures the scale of stimulation across patients.

#### PRC-ARC shift in WC synthetic data

The PRC-ARC shift is computed on WC synthetic data with phased-locked blocks of stimulation generated by the full model fitted to each patient. This time we can take full advantage of the model and compute bPRCs and bARCs from more trials than for patient data or model data in optimisation steps, and perform 10 repeats of 600 trials for the top 15 fits for each patient. The PRC-ARC shift is then measured as in section 2.1 for each of the 10 repeats, and shown in Figure 11. The large filled circles represent the mean of the 10 repeats for each patient fit. It appears that the PRC-ARC shift obtained for synthetic data of top patient fits mostly lie in the upper-left quadrant of the unit circle for all three patients 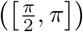, similarly to patient data. One fit to patient 6 is an outlier in terms of its shift, due to high effective stimulation (defined in the previous section). While the model can allow for a larger shift than 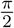, this is not the case for the linearised model, and the difference is the focus of the next section.

**Figure 10:**
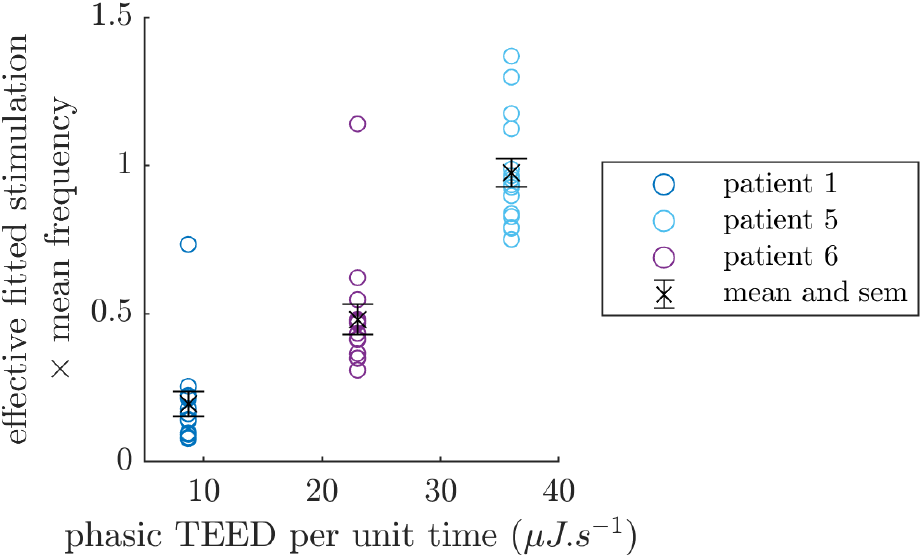
Fitted effective stimulation times mean frequency of *E* versus total electrical energy delivered per unit time by the device, for the three fitted patients. Showing the 15 best performing models for each patients, along with the mean and standard error of the mean error bars for each patient in black.

**Figure 11:**
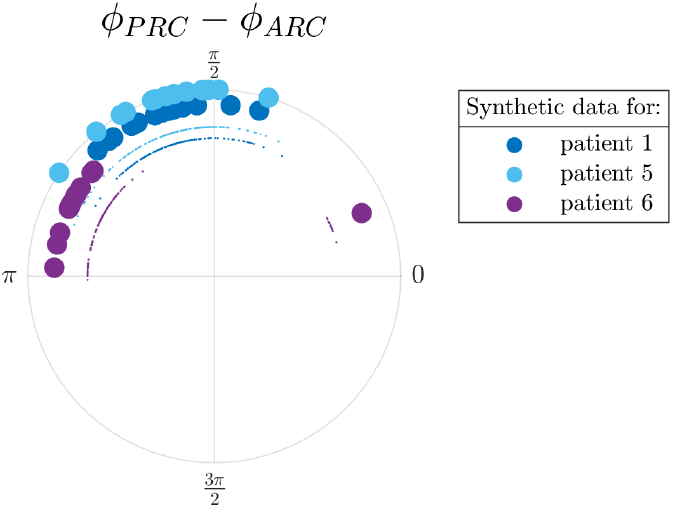
PRC-ARC shift in synthetic data (full WC model fitted to patients). For each patient, the shift for all 10 repeats of the top 15 fits is shown (smaller circles), as well as the repeat mean for each fit (larger circles). One repeat corresponds to 600 trials.

## 6 PRC-ARC shift in the model

The linearised model makes different predictions for patient fits than the full model, in particular in terms of PRC-ARC shift. The present section will look at the deterministic linearisation of patient fits, and then contrast it with the full model with noise.

### 6.1 Relationship between analytic response curves in the linearised fitted WC models

The first order PRC and ARC expressions derived in section 4 can be applied to the linearisation of the best WC models fitted to data from the three selected patients, and the Jacobians at the fixed points are

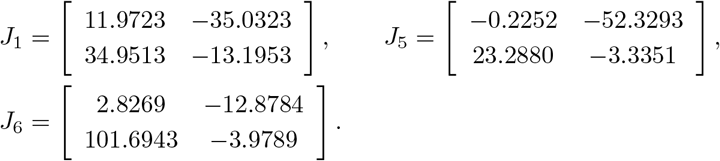

In the fits *b*_1_ = 0 or *b*_2_ = 0, which marginally simplifies equations (12) and (14). The curves obtained are shown in Figure 12. The same values as in section 4.7 are used for 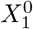 and *δX*_1_. Note that the stimulation delay Δ*t_stim_* is not shown – it affects both the PRC and the ARC and does not play a role in the PRC-ARC shift. More interestingly, we observe that *ω* ≫ |*σ*| in the 3 fits (see supplementary Table4 in Appendix I), suggesting that the response curves’ relationship described by equation (16) should approximately hold. This is indeed the case as shown in the third column of Figure 12. The decay is higher for patient 5 and as expected, the approximation is slightly worse for this patient (panel B3 in Figure 12). For small stimulation, the deterministic picture with patient parameters close to the fixed-point is that the PRC-ARC shift should be very close to 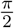. In what follows, we investigate the difference between this idealised picture and what is observed in synthetic data.

**Figure 12:**
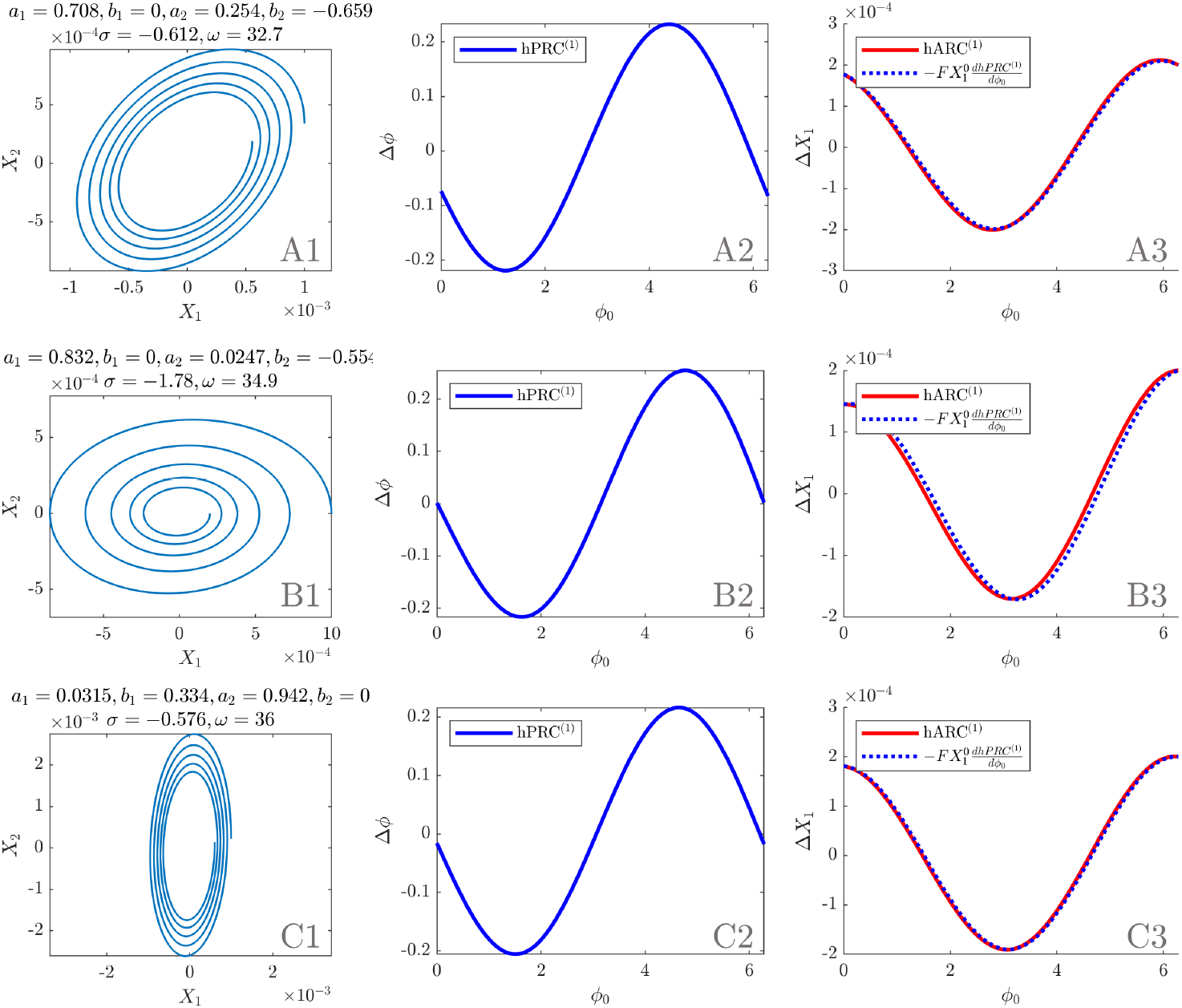
Analytical results for linearised patient fits (initial conditions as in the main text). First column: phase space. Second column: first order PRC as per equation (12). Third column: first order ARC as per equation (14) and opposite of the derivative of the first order PRC scaled by 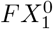. Panel A, B, and C correspond to patient 1, 5, and 6, respectively.

### 6.2 Accounting for the difference in shift between focus model analytic expressions and WC synthetic data

Four factors could account for the difference in PRC-ARC shift between the idealised picture given by analytic response curves with patient parameters (previous section) and what is observed in WC synthetic data (section 5.2). First, stimulation may be large enough that the Taylor expansions used to derive the analytic PRC and ARC expressions are no longer approximately valid. Second, tremor in patient fits may correspond to a regime not so close to the fixed point, compromising the linearisation validity. Third, the introduction of noise in the model may result in effects on the PRC-ARC shift that do not average out to zero. Fourth, in synthetic data, the response to stimulation is measured by the block method, which differs from the first order approach taken in our derivations. We next show that for the three best fits considered non-linearity is the main driver.

Ten repeats of 600 trials of synthetic data are generated for the linearisation of the best fits to each patient. The integration scheme with live phase tracking and stimulation is the same as described in section 5.1, only the stochastic differential equations are now

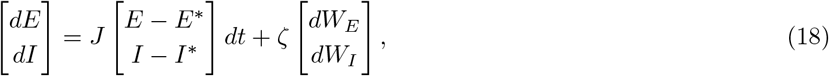

where *dW_E_* and *dW_I_* are Wiener processes, *ζ* the noise standard deviation (same values as in the non-linear case), *E*\* and *I*\* are the coordinates of the fixed point, and *J* is the Jacobian at the fixed point of the patient fit. The same values as in the non-linear case are used for the stimulation magnitude and delay, with the exception of patient 5, for whom the stimulation magnitude is set to a fifth of its value in the non-linear case, as higher values were seen to cause a breakdown of phase tracking, and result in unreliable response curves.

For each patient and for each of the 10 repeats, bPRCs and bARCs are obtained, and the PRC-ARC shift is then measured as in section 2.1. The results are shown in Figure 13 (middle), alongside the shifts measured from the response curves presented in section 6.1 (left), and the shifts measured in the full WC model (right). It can be seen that going from the analytic response curves to the linearised model (i.e. adding noise, measuring the response to stimulation via the block method and not a first order method, and using a finite stimulation magnitude rather than a infinitesimal stimulation), doesn’t affect the shift much. However, a substantial increase in the shift is obtained by introducing the non-linearity, which brings the shift in the upper-left quadrant, where patient data lie.

**Figure 13:**
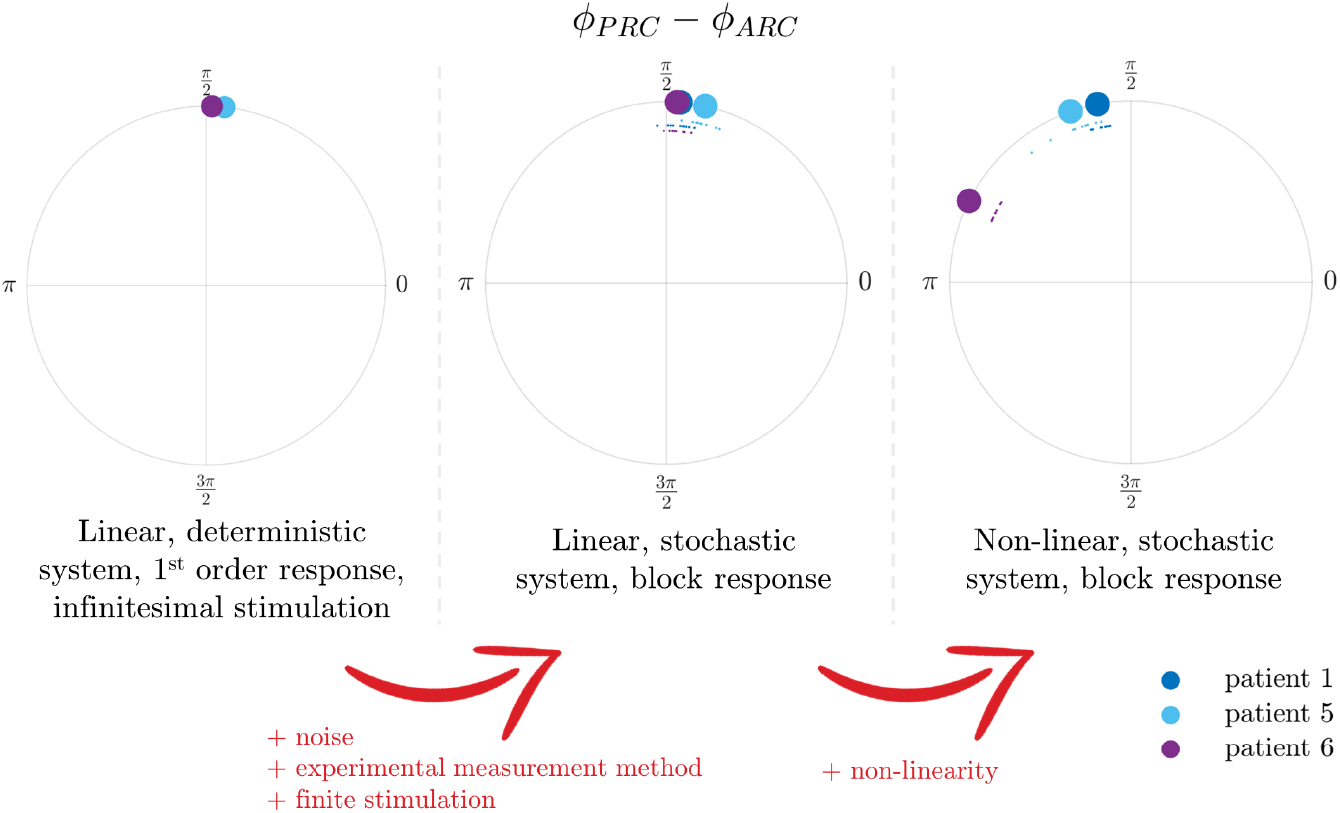
Non-linearity accounts for most of the difference in PRC-ARC shift seen in synthetic data (middle and right), when compared to the PRC-ARC shift derived in the focus model (left). When computed from synthetic data, the PRC-ARC shift of all 10 repeats is shown (smaller circles), as well as the repeat mean (larger circles). One repeat corresponds to 600 trials, only showing best fits for each patient.

## 7 Discussion

We showed that in a 2D linearised stable focus model, the first order PRC and the ARC are close to sinusoidal, in particular for small decay. Moreover, the PRC-ARC shift is close to 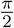. Half of the patients in our dataset had significant sinusoidal bPRCs and bARCs (an effect of stimulation phase could not be found in other patients in at least one of their response curves), and the significant patients have a PRC-ARC shift in the interval 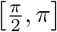. A full WC model can be fitted to tremor dynamics features and to the bPRC for these patients, and as hinted at by the similarities seen in the linearised focus model and the data, the best fits – a vast majority of stable foci – can reproduce the dependence of the effects of stimulation on the phase of stimulation. The best fits also reasonably predict the bARC, and notably what is approximately the best phase to stimulate. Compared to the 2D linearised focus, the non-linearities of the full WC model allow for a better reproduction of the phase dependence found in patient data, in particular as far as the PRC-ARC shift is concerned. Our full model can capture the behaviour of neural populations plausibly involved in the generation of tremor, which, together with its success in reproducing phase response and predicting amplitude response in patients, makes it a strong candidate for further study of phase-locked DBS.

### Phase definition

While asymptotic phase definitions are common in theoretical studies, experimental studies tend to favour instantaneous phase definitions such as the Hilbert phase. To reproduce the data, an instantaneous phase seems more appropriate than an asymptotic phase, as there is no indication of stimulation happening on or close to an attractor. It has been shown recently in [36] how an operational definition of the phase can describe transient spiking, when an asymptotic phase does not capture the phase dependence of transients. In this study, our phase definition is the Hilbert phase of the tremor data. It is therefore referenced to the maximum of the tremor oscillations (represented by the first coordinate of the dynamical system), and does not require a limit cycle. The Hilbert phase is an angle in the analytic signal space, it does not generally grow linearly with time, and is a protophase [37]. This is not a concern from the perspective of describing patient data, as this is the observable choice we are making for both the data and the model. Commonly used with data, the Hilbert transform has also been proposed as a robust method to measure steady state PRCs in single neuron models [38]. Moreover, stimulation is assumed to be small in our analytical expressions (section 4), but not in the full model, contrary to standard asymptotic phase reduction strategies.

### Linearisation

The response curves derived for the linearisation of a 2D focus in section 4 can be related to previously published expressions. In particular, the infinitesimal PRC for radial isochron clocks has been derived in [39], and has been recently included in [40] under the larger umbrella of general radial isochron clocks. The radial clock case (*K*(*ϕ*) = *ω* in [40]) perturbed along the first dimension agrees with our equation (12) for the case of a circular flow (see section 4.7). For this simple system, the asymptotic phase response is the same as the first order Hilbert phase response.

Moreover, for small decay, the best phase to stimulate corresponds to the maximum positive slope of the first order PRC in the response curves derived. In fact, the first order ARC is simply a scaled version of the opposite of the first order PRC derivative. A similar relationship has been first reported in a theoretical study in the context of an individual oscillator [41], and more recently in [15] in the context of population response curves of a Kuramoto model. It is noteworthy that we found a similar result for the linearisation of any 2D focus (i.e. any model whose dynamics obey equation (4)) with slow decay, and in particular for the linearisation of the WC model, another popular neuroscience model very different in essence from coupled oscillator models. In the thermodynamic limit and under certain assumptions about the distribution of oscillator frequencies, the Kuramoto model can be reduced to a two-dimensional system [42]. Our results are applicable to the linearisation of a fully desynchronised reduced Kuramoto model observed through *X*_1_ = *ρ* cos *θ* where **r** =*ρe^iθ^* is the order parameter (*ρ* is the modulus and *θ* the angle in the complex plane). Such a system therefore satisfies equation (16) as well (for small decay).

Our derivations do not assume proximity to a limit cycle, and this allows the study of the dependence of the response to stimulation on the amplitude of the oscillations for a given model (limit cycles do not have an amplitude variable in the case of infinitesimal perturbations). In the linearisation, the PRC is found to be inversely proportional to the amplitude of the oscillations before stimulation (see equation (12)), while the ARC does not dependent on it.

Because the block method phase and amplitude response used in the rest of paper are normalised by the number of pulses and blocks are only about 25 period long, it seems legitimate to think that, although they are different objects, the first order response to a single pulse (hPRC^(1)^ and hARC^(1)^) and the block method response (bPRC and bARC) could be related, and in particular that they might share similar PRC-ARC relationships. Part of the connection hinges on our proof that the phase definition in the linearisation of the focus model overlaps with the Hilbert phase when the decay is small compared to the rotation (section 4.2). And indeed, the PRC-ARC shift predicted by our expressions derived for the first order response to one pulse of stimulation in a linearised focus is very close to the shift obtained by the block method on linearised WC synthetic data (see Figure 13). Our analytical derivations provide a rationale to fit the full WC model to data and an intuition for why the model can predict patient ARC, but do not offer an exact analytic treatment of the block method. Specifically, individual pulses in a block may have different effects depending on where they are located in the block and depending on stimulation history within the block [11].

### Fitting procedure

Fits were performed using the generalized pattern search algorithm on many sets of random initial parameters. This approach was chosen for its robustness and computational efficiency in a non-smooth, non-convex landscape with four non-linear features and 10 parameters, despite requiring the use of a supercomputer. In particular it has been deemed superior in finding better fits to the simplex algorithm. The implementation used also has the additional benefit of being able to handle failed simulations (which occasionally happen as response curves with 12 phase bins can not be obtained for some parameters). However the fitting procedure results in many “good” local optima. What these “good” sets of parameters have in common and what they can tell us about the patients we are fitting to is not easily addressed with our current fitting strategy. Even real biological networks may have redundancies, and may exhibit the same behavior under different network configurations. Approximate Bayesian computation [43, 44] allow to approximate the posterior distribution over parameters for intractable likelihoods, hence to answer the question what is the space of parameters consistent with the data. Whether approximate Bayesian computation methods could successfully tackle a complicated landscape and provide more meaningful insight on fitted model parameters in the setting of the present work is a promising avenue for further research. A limitation of our fitting method is related to the integration scheme: to reduce computation cost, the Euler step used in the first optimisation process is 1 ms. The top 20 best fits are then re-optimised based a Euler step of 0.1 ms, and results are produced with this finer time step, as dynamics can be qualitatively different (further reduction in the Euler step has not been seen to change the dynamics). While the necessities of phase-locked stimulation precludes the use of built-in, powerful integration schemes, a more advanced event-based stochastic integration scheme could remove the need for a second optimisation while keeping the computation cost down. The performance of our simple phase-tracking strategy is good for patient 1 and 6 and satisfactory for patient 5 (see Figure 16 in Appendix). Response curves are obtained based on the actual Hilbert phase of stimulation in a post-hoc manner, which makes up for the reduced performance observed for patient 5. Still, more accurate algorithms could be explored. Our zero-crossing strategy would benefit from a better frequency estimate for the current period (currently simply the period of the previous period) and more robustness to noise. The proposition of [45] involving autoregressive forward prediction and the Hilbert transform is attractive, although its computational cost may be high, and some parameters need to be adjusted for each time series.

### Non-linear WC model

The full WC model is fitted to data with Gaussian white noise (equation (1)). The best performing fits are stable foci for all three patients, and very few limit cycles are found in the top 15 fits for all three patients. One is found for patient 1 (shares the 1st place with a stable focus - distance between mean costs only 30% of the standard error of the mean), one for patient 5, and none for patient 6. In the stable focus regime, noise brings the system away from the stable fixed point, and the interaction of the noise with the dynamics of the system makes the reproduction of patient tremor possible. In the absence of noise, the system would converge to the stable fixed point and no tremor would be generated, so symptoms are related to the noise level in this model. Instead of noise, tremor-like activity may be obtained by exploiting chaotic dynamics arising from coupling several WC models together [46], but this would significantly increase the complexity of the model (more on increasing complexity in the last part of this section).

In fitting our thalamic model to tremor acceleration, we are assuming thalamic activity and tremor are directly related as mentioned before (see section 5.1). Tremor activity is however expected to lag thalamic activity due to conduction delays. The accelerometer used to measure tremor is also expected to introduce an electromechanical coupling delay. In the model, we allow for a stimulation delay Δ*t_stim_* between the stimulation trigger and the time when stimulation is actually delivered to the excitatory population. This parameter is fitted to the data, and gives the model the ability to shift its bPRC in phase. Fitted stimulation delays are hundreds of milliseconds, and conduction and accelerometer delays (tens of milliseconds) only account for a small part. The higher fitted values are required by the model to match data bPRCs. With our candidate VIM/nRT mapping in mind, the higher fitted values remain unexplained on the biology side, although as mentioned before tremor generation and ET DBS are not fully understood. It is interesting to note that the stimulation delay of the best performing model for patient 5 is longer than one period (see Table 2). This is found consistently in the top three best fits, and reducing the delay to its value modulo the average period substantially reduces the quality of the bPRC fit. Besides this short term delay, our model does not include medium or long term plasticity effects, which are not expected to be strongly present in the recordings as stimulation is only delivered for periods of 5 seconds in a row. In our model, stimulation is provided to the *E* population via a direct increase in the population activity. While stimulation is provided via the sigmoid function of the excitatory population in other studies [18], we found this approach too restrictive due to sigmoid saturation, and inadequate to reproduce the full extent of the response to phase-locked DBS in some patients. As a reminder, the choice of stimulating the excitatory population rather than the inhibitory population is made for biological consistency, as the VIM is the most common stimulation target in ET DBS.

The success of the WC model in predicting patient ARCs when fitted to their PRCs is partially explained by its ability to modulate the PRC-ARC shift. The PRC-ARC shift in the full model can reach the range found in patients while the linearised version of the WC is limited to the close vicinity of 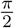. The response curves of the full WC model are also better at reproducing the data and can vary from pure sinusoids. However there is still some room for improvement in reproducing the shift, in particular as far as patient 1 is concerned (patient shift quite a bit larger than the model). The model can allow for a larger shift as shown by a fit hand-picked in the top 15 shown in Figure 17 in Appendix H. The PRC-ARC shift could be selected as an additional feature to fit to to improve ARC reproduction.

In its 2 population version, the suggested mapping of the excitatory and inhibitory populations (VIM and nRT) is not the only possibility. Other candidates include antidromically stimulated structures at the cerebellar level or below, such as DCN as the inhibitory population, and the inferior olive as the excitatory population. The model could also be extended by including more populations. With our current mapping in mind, the cortex and the DCN could be turned into populations of their own, which would make the model four dimensional. As suggested in [18], the inferior olive which provides input to the DCN could also be modelled, and the spatial extent of the VIM could be accounted for by splitting it in two populations or more. Increasing the number of populations would however increase the number of parameters of the model, and make the optimisation process more computationally intensive, and the model more prone to over-fitting. In contrast, the incorporation of additional loops in the model architecture may help explain the inertia in stimulation effects discussed above. Nevertheless, the model seems to be able to reproduce the data in its current state, which suggests an increase in complexity is not warranted. It is remarkable that one excitatory/inhibitory loop seems to be enough to model the phase-dependent effects of ET DBS. It gives some support to the hypothesis that sub-circuits of the central tremor network may behave as individual oscillators entraining each other [47].

## 8 Conclusion

The focus WC model with noise can be fitted to ET patients with both response curves showing significant phase dependence. The model reproduces the phase dependence of the response to stimulation as well as predicts the amplitude response to stimulation, which directly relates to tremor reduction. Phase-locked stimulation promises less stimulation, hence less side effects for the same clinical benefits, which would be highly desirable for patients. Our study positions the WC as a strong candidate to model the effects of phase-locked DBS. Its ability to describe all patients with both response curves significant in at least one of our tests should be re-assessed as more data becomes available, both in terms of number of patients and recording length. Phase dependent activity is thought to play a central role in physiological information processing [48, 49], and in our analytical derivations, the phase of the linearised model was defined in a way that does not depend on modelling oscillations by a limit cycle, and that for small decay overlaps with the Hilbert phase, which is widely used in experiments. Finally, as far as ET generation is concerned, we showed that a single excitatory/inhibitory loop is enough to reproduce both the dynamics of the tremor and the phase dependent effects of stimulation, however it should be non-linear.

## List of abbreviations

ARC: amplitude response curve
DBS: deep brain stimulation
DCN: deep cerebellar nuclei
ET: essential tremor
FDR: false discovery rate
nRT: reticular nucleus
PD: Parkinson’s disease
PDF: probability density function
PRC: phase response curve
PSD: power spectrum density
TEED: total electrical energy delivered
VIM: ventral intermediate nucleus
WC: Wilson-Cowan

## Appendices

We include here technicalities on approximating the Hilbert phase in the linearisation (Appendix A), details of the derivations leading to response curves analytical expressions in the linearised system (Appendices B to D), and the procedure used to obtain WC parameters from a given Jacobian (Appendix E). We also present details of the two-step optimisation used for fitting to patient data (Appendix F), the implementation of live-phase tracking and stimulation (Appendix G), as well as supplementary figures (Appendix H) and supplementary tables (Appendix I).

## A Hilbert transforms of sine and cosine exponential decays with error terms

The goal here is to show that 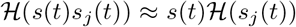 for *j* = *c, n*, with *s*(*t*) = *e^σ|*t*|^, s_c_*(*t*) = cos *ωt*, and *s_n_*(*t*) = sin *ωt*. The Bedrosian identity [35] states that the Hilbert transform of the product of a low-pass and a high-pass signal with non-overlapping spectra is the product of the low-pass signal and the Hilbert transform of the high-pass signal. The spectrum support of *s* is ℝ, but for low decay compared to the rotation, the spectrum of *s* is very small where it overlaps with the spectra of *s_c_* or *s_n_*. The equality given by the Bedrosian identity turns into an approximation, and inspired by the proof in [35], we can calculate error terms. Let *S* and *S_c_* be the Fourier transforms of *s* and *s_c_* respectively:

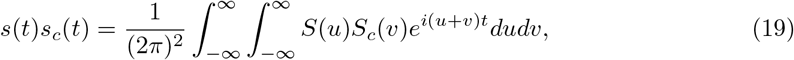

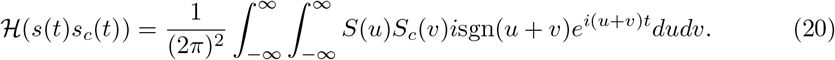

The Fourier transform of *s_c_* is given by *S_c_*(*v*) = *π*[δ(*v* − *ω*) + *δ*(*v* + *ω*)], so

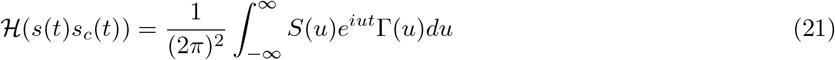

where 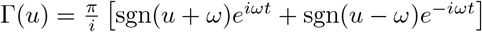. This can be simplified as

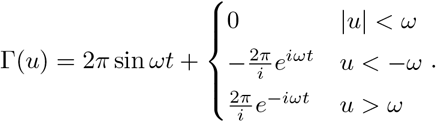

The Fourier transform 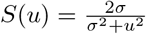 is even, therefore

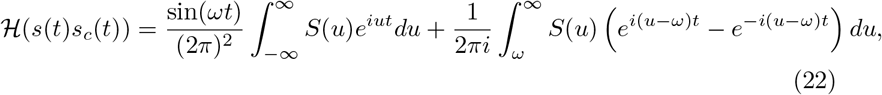

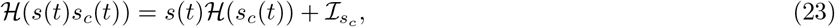

with

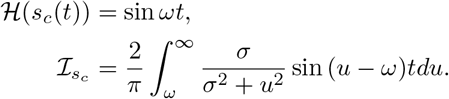

A similar derivation provides

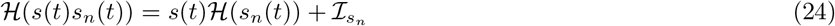

with

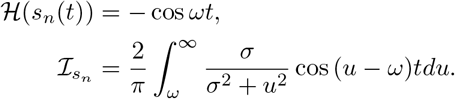

Numerical integration proves that for *ω* ≫ |*σ*|, and in particular in the case of the patients we are interested in, 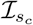 and 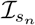 are under 5% of the signal scale for about 12 periods (see Figure 14). This is more than enough for our purposes as only one period is needed to derive response curves. It is therefore reasonable to ignore 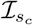 and 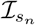.

**Figure 14:**
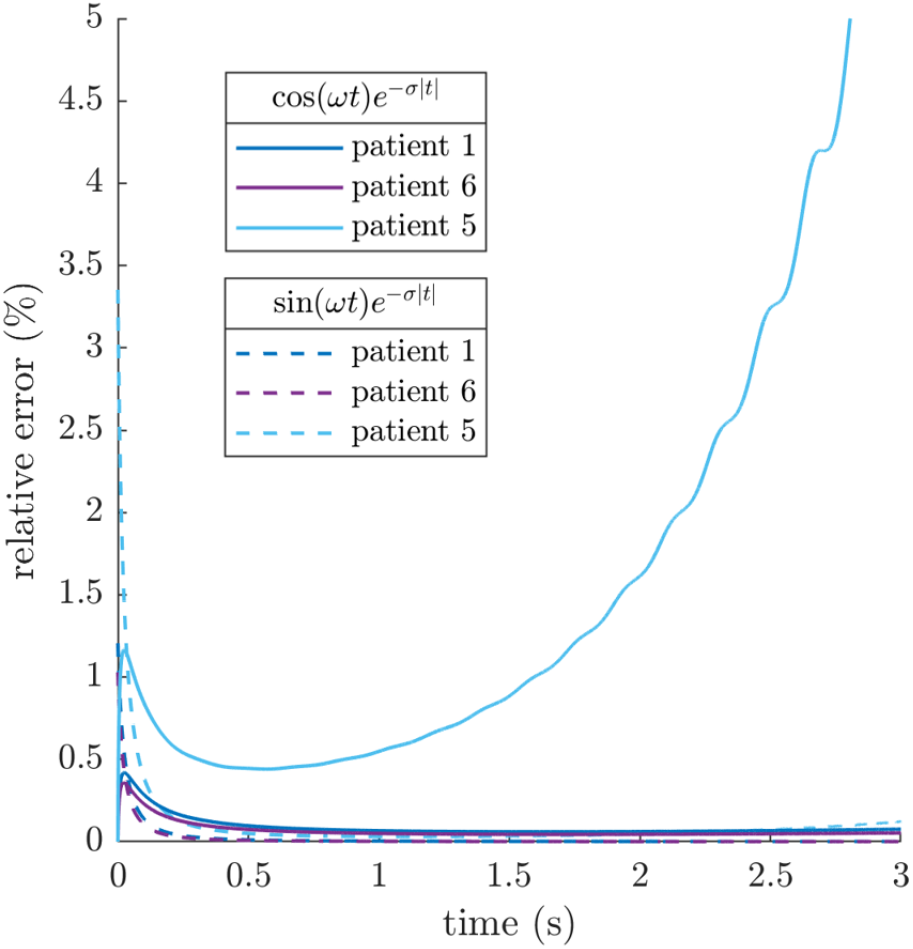
Relative error made across patients in estimating 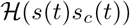 by 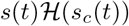 (solid lines) and 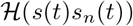 by 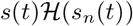 (dashed lines). The error is calculated as the ratio of 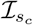 (respectively 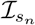) over the modulus of the numerical Hilbert transform of the signal, which is the envelope of the signal. The relative error is under 5% in all cases for at least 12 periods.

## B Reference trajectory without stimulation

Let us find the coefficients *K_ref_* and 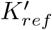 of the trajectory starting at *t* = 0 at a maximum of the first coordinate 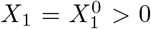. With the choice *ϕ* = *ωt*, this will ensure we are referencing the phase to the maximum of *X*_1_. It should be noted at this point that we are not using the nullcline equations in what follows as we are interested in the dependence of the response on the rotation *ω* and the decay σ. From the initial condition at *t* = 0,

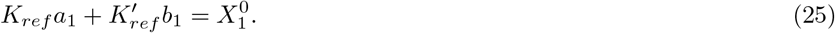

Additionally, 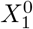 being a maximum requires that 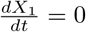 at *t* = 0, therefore

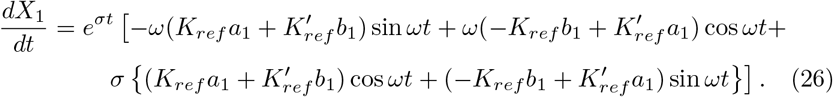

Using the condition at *t* = 0,

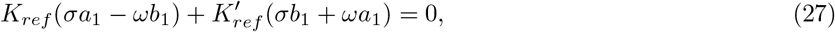

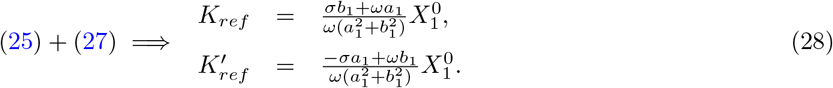

We are excluding the case where the denominator in (28) is equal to zero, which corresponds to both *a*_1_ and *b*_1_ being zero, which would imply *X*_1_(*t*) = 0. Also note that by picking a positive *X*^0^_1_, we are ensuring that the null derivative corresponds to a maximum of *X*_1_ rather than a minimum.

## C Trajectory with stimulation

Let us determine what the coefficients *K_stim_* and 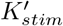 are for the stimulated trajectory (still constrained by the dynamics of equation (4)). We have

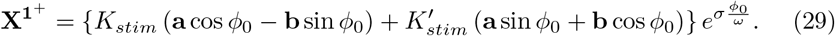

Solving for *K_stim_* gives

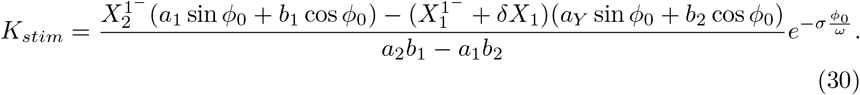

Plugging in 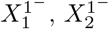, and the expressions for *K_ref_* and 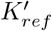 yields

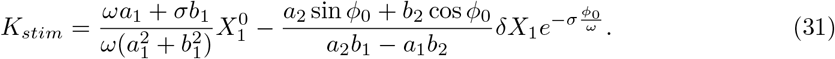

Similarly for 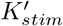, using the previous result:

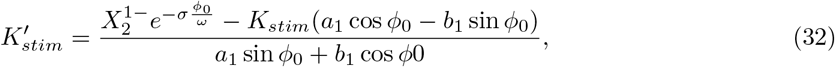

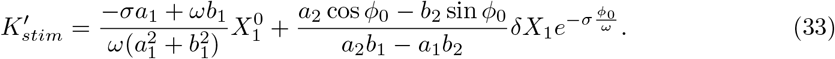

## D Phase at the next maximum of *X*_1_ on the stimulated trajectory

We are looking for *ϕ_max_* such that 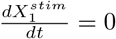 at *ωt* = *ϕ_max_*. This give us

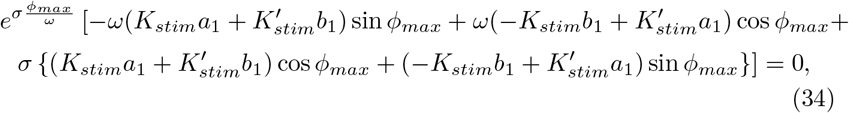

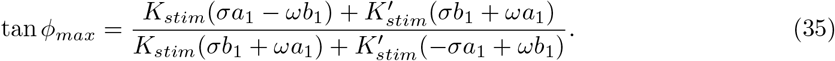

The phase *ϕ_max_* is returned by the arctan function in 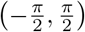, and corresponds to the previous peak on the stimulated trajectory extended backwards. The next peak has the same phase (mod 2*π*) as the expression in square brackets in equation (34) is 2*π*-periodic.

## E Finding WC parameters corresponding to a given Jacobian

The Jacobian of (1) evaluated at (*E*\*, *I*\*) can be simplified by making use of *f*′(*x*) = *βf*(*x*)(1 − *f*(*x*)). We also have

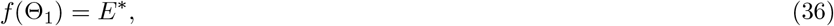

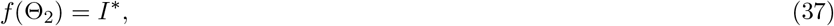

with

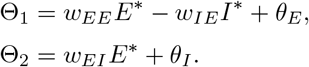

The Jacobian of (1) evaluated at (*E*\*, *I*\*) is therefore given by

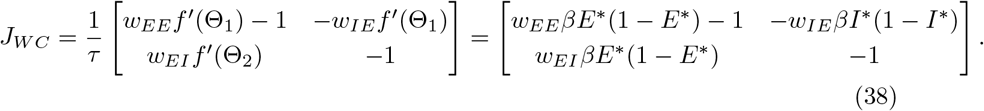

We are interested in finding WC parameters so that the linearisation of the WC model at the fixed point will be characterised by a given Jacobian matrix

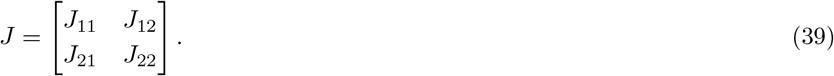

If we pick values for *β*, *E*\* and *I*\*, the remaining parameters can be obtained by equating (38) and (39), and by re-arranging equations (36) and (37). Parameters in supplementary Table 3 were obtained using this method, which yields

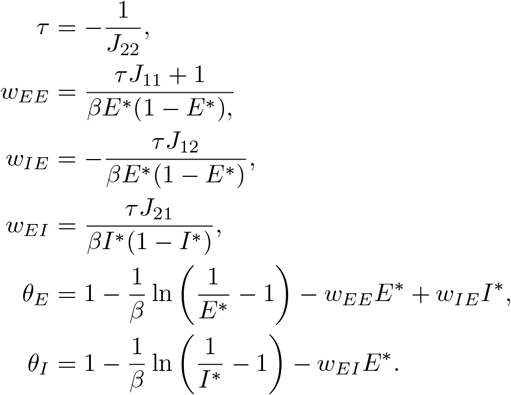

## F Two-step optimisation

The optimisation procedure is as follows. For each patient, random sets of parameters are picked from uniform distributions (bounds in supplementary Table 5). To improve the efficiency of the optimisation, we accept parameters only if the PSD peak of the corresponding model (without stimulation) is within 1 Hz and 25% in magnitude of the data PSD peak. Once 2500 parameters have been accepted, we put them through local optimisations. Local optimisations are carried out using a direct search algorithm called the generalized pattern search algorithm. Parameters are put on a similar scale to improve search robustness, and hard limits are given to the optimiser (see supplementary Table 5 in Appendix I). Optimisations are performed in parallel on a supercomputer. A time step of 1 ms is used for the fits (a period is about 200 ms). At the end of this process, the 20 best performing sets of parameters were put through more local optimisations with a finer time step of 0.1 ms and stop criteria leaving room for more steps. The finer time step is also used to produce the results shown in section 5.2).

The implementation of the generalized pattern search algorithm used is Matlab’s patternsearch optimiser with the poll method “positive basis 2N” and the following stop criteria:

- main optimisation (time step of 1 ms): mesh size of 10^-4^, function call budget of 800,
- second optimisation (time step of 0.1 ms): mesh size of 10^-5^, function call budget of 1000.

## G Live phase tracking and stimulation

One simulation consists of 600 trials with 12 blocks of phase-locked stimulation each. As in the experimental paradigm, blocks last 25s, and inter-block intervals are 1s. Inter-trial intervals are 5s, and the first trial starts after about 200 periods. During this initial time, the mean of *E* and the standard deviation of *E*, *σ_sim_*, are obtained from about 20 periods after a ramp-up of about 40 periods. Phase-tracking subsequently starts: *E* is centered and a threshold *T* = 0.2*σ_sim_* is used to track positive zero-crossings. The use of hysteresis via a threshold was found critical to handle the noise included in the model. We define a positive zero-crossing as happening when

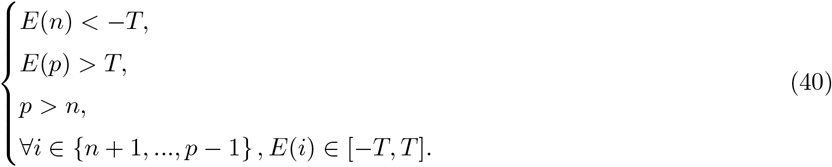

These conditions are constantly monitored, and if found true, a positive zero-crossing is declared to have happened at time step 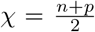. We evolve the zero-crossing phase according to a frequency based on the previous period, and if *χ_k_* is the last positive zero-crossing to have occurred, the current value of the zero-crossing phase is given by

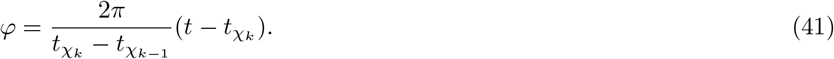

If the value of 2*π* is reached, the phase value is set to 0 until the next positive zero-crossing is detected. Stimulation is provided after *φ* reaches the target phase for the block, and the stimulation trigger is recorded Δ*t_stim_* before stimulation occurs. If the zero-crossing phase hasn’t reached the target stimulation phase yet when the next positive zero-crossing is detected, stimulation is provided right then. As in [11], a pulse of stimulation consists of six quick bursts at 130 Hz.

## H Supplementary figures

**Figure 15:**
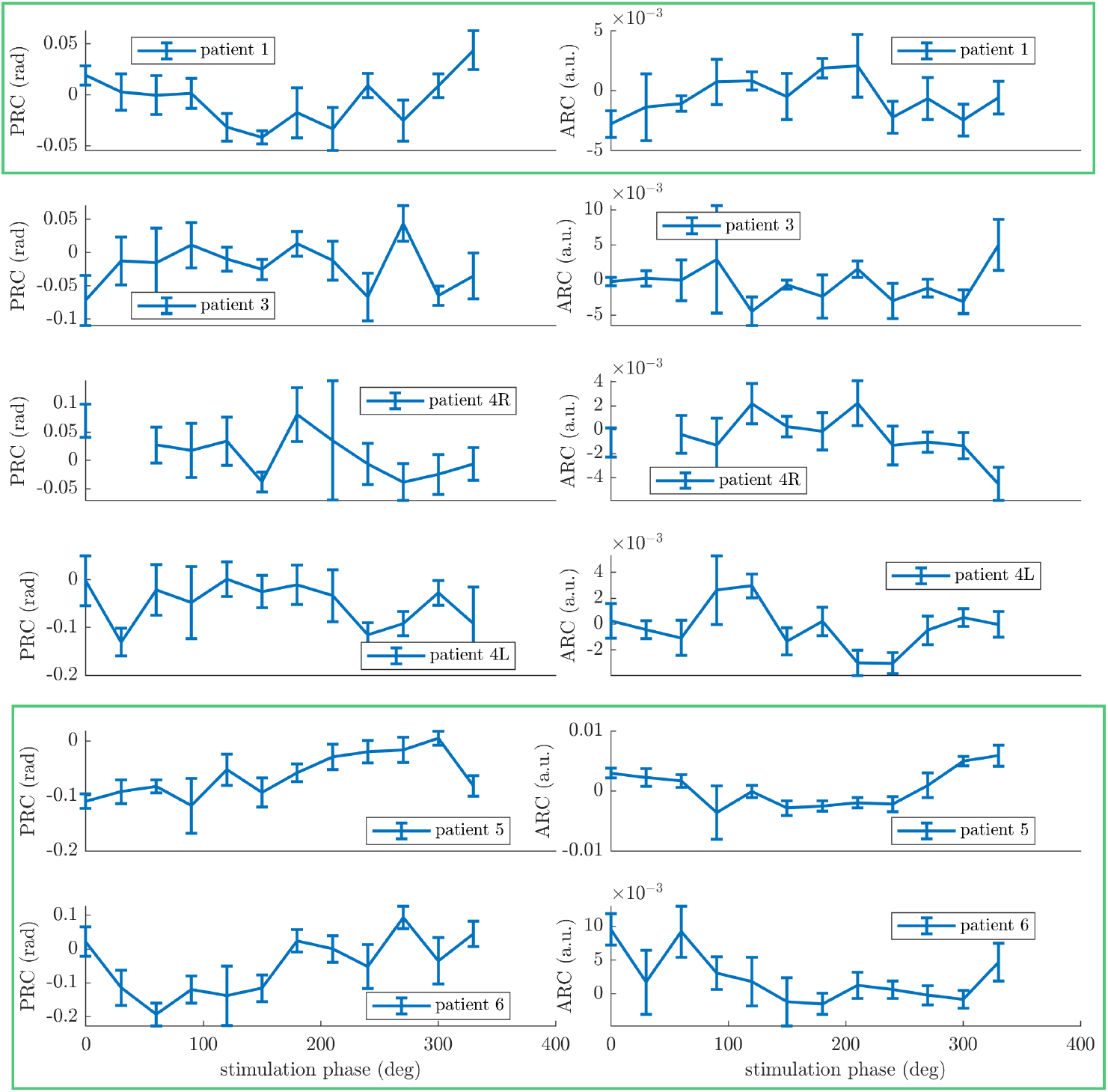
Patients’ PRCs (first column) and ARCs (second column) obtained as described in section 2.1. Datasets with both response curves significant according to at least one of our statistical tests under FDR control are highlighted with green rectangles.

**Figure 16:**
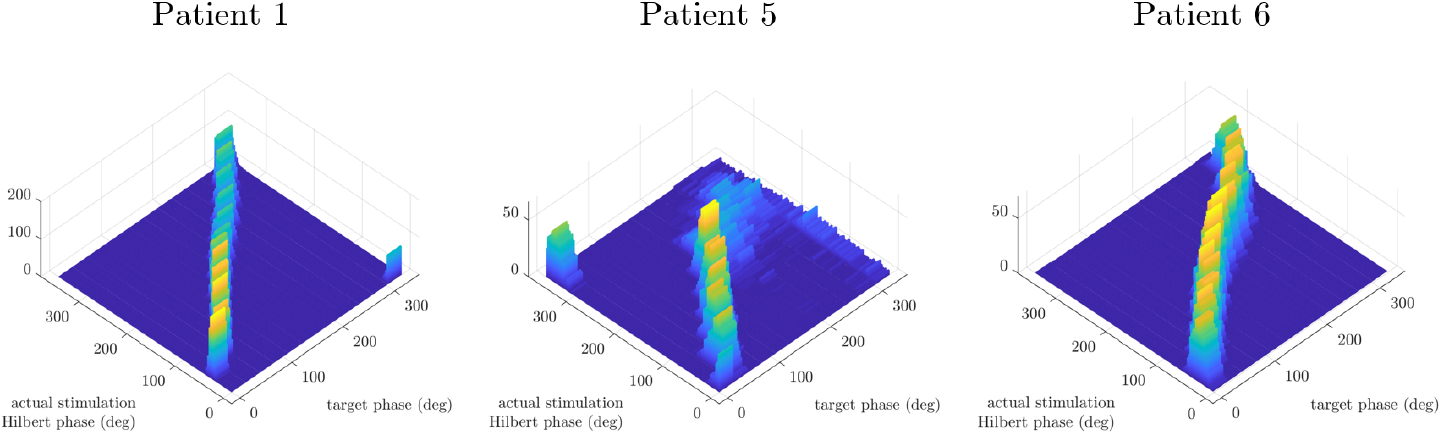
Phase tracking illustrated in the three fitted patients by histograms of the pair (target stimulation phase for the stimulation block, average of actual Hilbert phase at stimulation for the stimulation block). The actual Hilbert phase is obtained post-hoc after filtering. A block average includes averaging across bursts and within the block. Averages are obtained using circular means. The effect of the stimulation delay was removed, and phases are reference to positive zero-crossings. Phase tracking is satisfactory for all patients, although tracking is less precise for later phases in patient 5.

**Figure 17:**
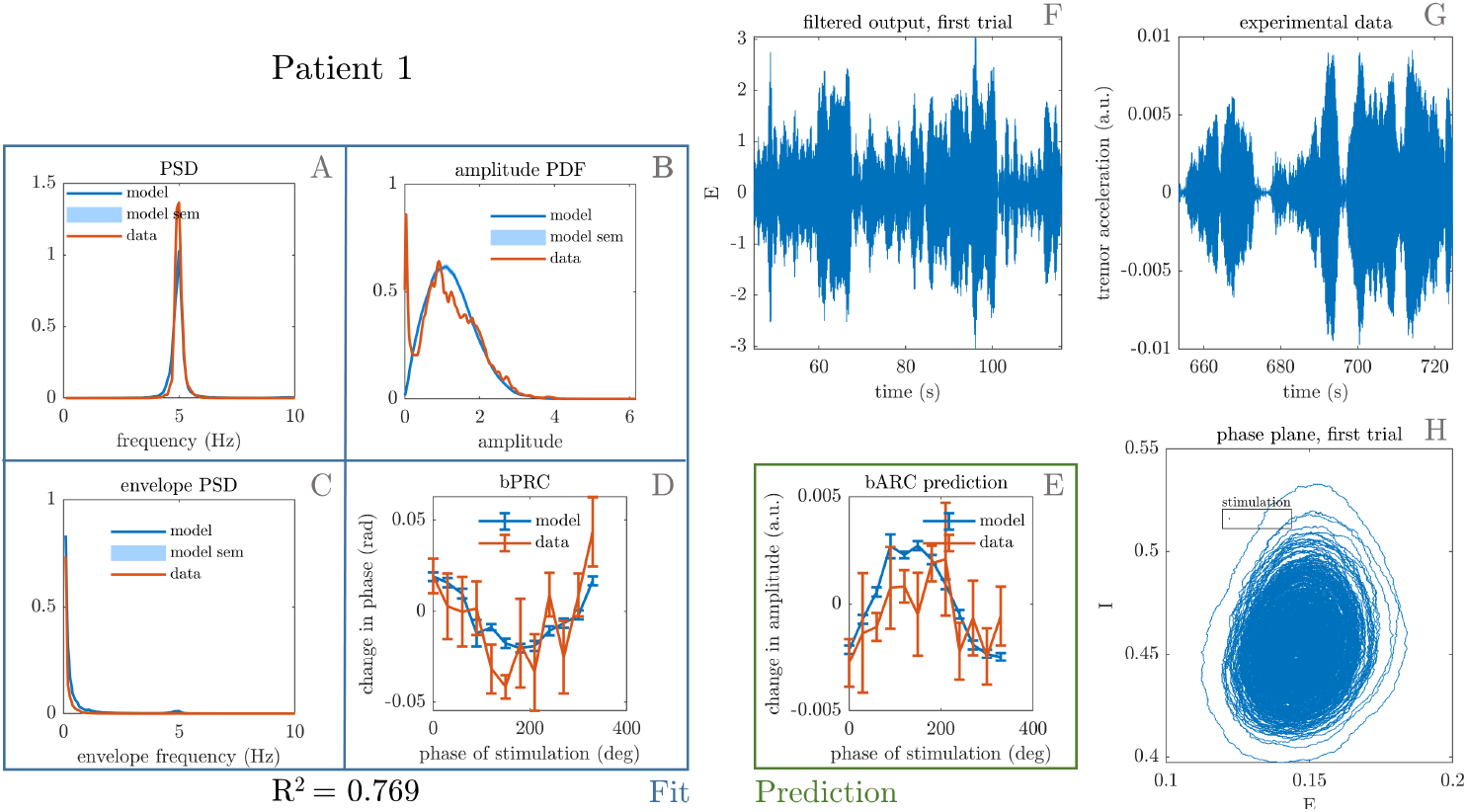
Fit to patient 1 showing the best PRC-ARC shift. The four features that were included in the cost function are shown on the left, namely tremor PSD (A), tremor envelope PDF (B), tremor envelope PSD (C) and PRC (D). The model better predicts the data ARC (E) thanks to a PRC-ARC shift closer to that of the data. The model phase plane is shown in H, and the model tremor time-series (F) is shown next to the patient tremor time series (G). The framed black bar in H indicates the fitted stimulation magnitude to scale.

## I Supplementary tables

**Table 3:**
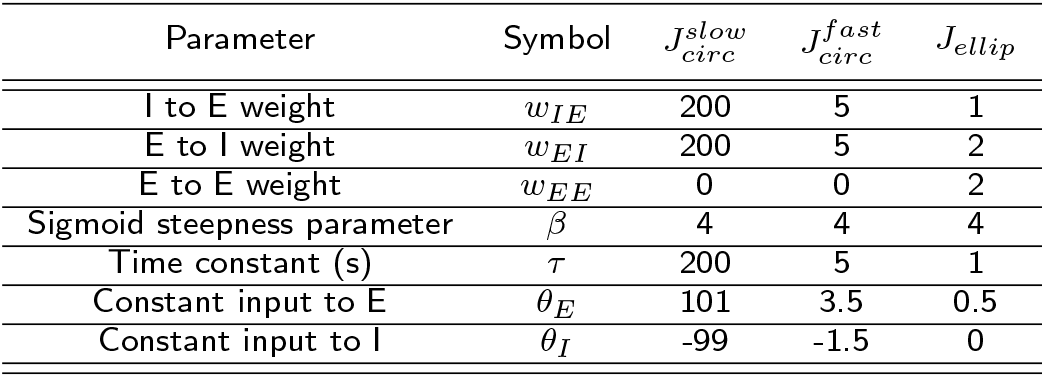
WC parameters corresponding to the Jacobians presented in section4.7. The steepness parameter *β* was set to 4, *E*\* and *I*\* to 0.5, and parameters were determined according the method presented in Appendix E.

**Table 4:**
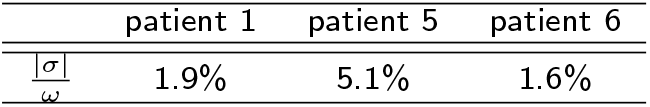
|*σ*|/*ω* ratios in the linearisation of patient fits.

**Table 5:** Lower and upper bounds of parameters uniform distributions used to generate initial parameters for fitting, and hard limits enforced by patternsearch during the optimization process.

| Parameter | Symbol | Initial parameter distribution |  | Hard limits enforced by optimizer |  |
| --- | --- | --- | --- | --- | --- |
|  |  | Lower bound | Upper bound | Lower bound | Upper bound |
| I to E weight | $w_{IE}$ | 0 | 10 | 0 | 30 |
| E to I weight | $w_{EI}$ | 0 | 10 | 0 | 30 |
| E to E weight | $w_{EE}$ | 0 | 10 | 0 | 30 |
| Sigmoid steepness parameter | $\beta$ | 0 | 10 | 0 | 30 |
| Time constant (s) | $\tau$ | 0 | 0.3 | 0 | 0.5 |
| Constant input to E | $\theta_E$ | -2 | 10 | -30 | 30 |
| Constant input to I | $\theta_I$ | -10 | 2 | -30 | 30 |
| Noise standard deviation | $\zeta$ | 0 | 0.1 | 0 | 0.3 |
| Stimulation magnitude | $\delta E$ | 0 | 0.02 | 0 | 0.1 |
| Stimulation delay (ms) | $\Delta t_{stim}$ | 0 | 250 | 0 | 500 |

## Acknowledgements

The authors would also like to acknowledge the use of the University of Oxford Advanced Research Computing (ARC) facility in carrying out this work http://dx.doi.org/10.5281/zenodo.22558

## Availability of data and material

The datasets analysed in the current study are available online [30].

## Author’s contributions

BD was involved in the conceptualization of the study, carried out the statistical, analytical and computational studies, and wrote the paper. GW provided guidance for the optimisation work. CB and RB provided supervision and were involved in the conceptualization of the study. HC and PB collected the original tremor data. All authors were involved in the discussion of the results and edited the paper.

## Ethics approval and consent to participate

Not applicable

## Consent for publication

Not applicable

